# H_2_S remodels mitochondrial ultrastructure and destabilizes respiratory supercomplexes

**DOI:** 10.1101/2024.10.30.621162

**Authors:** David A. Hanna, Brandon Chen, Yatrik M. Shah, Oleh Khalimonchuk, Brian Cunniff, Ruma Banerjee

## Abstract

Mitochondrial form and function are intimately interconnected, responding to cellular stresses and changes in energy demand. Hydrogen sulfide, a product of amino acid metabolism, has dual roles as an electron transport chain substrate and complex IV (CIV) inhibitor, leading to a reductive shift, which has pleiotropic metabolic consequences. Luminal sulfide concentration in colon is high due to microbial activity, and in this study, we demonstrate that chronic sulfide exposure of colonocyte-derived cells leads to lower Mic60 and Mic19 expression that is correlated with a profound loss of cristae and lower mitochondrial networking. Sulfide-induced depolarization of the inner mitochondrial membrane activates Oma1-dependent cleavage of Opa1 and is associated with a profound loss of CI and CIV activities associated with respirasomes. Our study reveals a potential role for sulfide as an endogenous modulator of mitochondrial dynamics and suggests that this regulation is corrupted in hereditary or acquired diseases associated with elevated sulfide.

**Significance Statement:** Hydrogen sulfide is a product of host as well as gut microbial metabolism and has the dual capacity for activating respiration as a substrate, and inhibiting it at the level of complex IV. In this study, we report that chronic albeit low-level sulfide exposure elicits profound changes in mitochondrial architecture in cultured human cells. Disruption of mitochondrial networks is reversed upon removal of sulfide from the growth chamber atmosphere. Sulfide-dependent depolarization of the inner mitochondrial membrane is associated with loss of cristae and respiratory supercomplexes. Our study reveals the potential for sulfide to be an endogenous regulator of mitochondrial ultrastructure and function via modulation of electron flux and for this process to be corrupted in sulfide dysregulated diseases.

## Introduction

Mitochondria are central hubs of energy metabolism that support oxidative phosphorylation via proton-coupled electron transfer through complexes in the electron transport chain (ETC), which are located in cristae. The latter are dynamic sac-like compartments formed by the inner mitochondrial membrane whose shape changes in response to physiological conditions (1). Arrays of dimeric ATP synthase induce curvature at cristae ridges while the proton-pumping complexes I, III and IV (referred to as CI, CIII and CIV) but not CII, are organized in supercomplexes or respirasomes of varying stoichiometry on cristae flanks (2). Cristae dynamics influence respiratory supercomplex stability and function and cristae disruption is associated with supercomplex destabilization (3). While the physiological relevance of supercomplexes remains elusive, roles such as protection against excessive reactive oxygen species (ROS) (4) formation, tuning electron flux for optimal substrate utilization (5), and boosting efficiency via proximity effects (6, 7), have been reported.

Cristae junction architecture is maintained by MICOS (mitochondrial contact site and cristae organizing system), the GTPase Opa1 (optic atrophy-1) and other proteins (1). The constitutive YME1L, and the stress-responsive Oma1 proteases cleave Opa1 into various short isoforms. A range of stimuli, including oxidants, ATP depletion, and dissipation of the transmembrane potential activate Oma1, influencing mitochondrial networking and connectivity (8, 9), highlighting the interconnection between cristae morphology and organelle dynamics (10). Mic19, a key stabilizing component of MICOS, and the oligomeric stability of the metalloprotease Oma1, are both sensitive to redox changes (11, 12). Regulation of mitochondrial form and function by the transmembrane potential and by changes in the ambient redox state, is a strategy for responding to changes in energy demand.

Hydrogen sulfide (H_2_S) is a potential candidate for modulating mitochondrial form and function, via its dual ability to interact directly with the ETC and its role as a signaling molecule (13–16). H_2_S is synthesized in the cytosol and in mitochondria (17, 18), and readily permeates membranes (19). ER stress, amino acid restriction, and hypoxia increase H_2_S synthesis (20–23). At low concentrations, H_2_S serves as an ETC substrate, reducing CoQ, as it is itself oxidized to glutathione persulfide by sulfide quinone oxidoreductase (SQOR) (24) (Fig. 1A), and potentially, by reducing cytochrome c (25). At higher concentrations, H_2_S inhibits complex IV, leading to respiratory inhibition that is long-lived (26), which in turn, induces a reductive shift in redox cofactors like NAD^+^ and CoQ (27, 28) and increases aerobic glycolysis (29) and lipid biogenesis (30). Inhibition of forward electron transfer by sulfide is predicted to ripple out into the intermembrane space, impairing the Mia40-dependent oxidative protein folding pathway, which uses cytochrome c as an electron acceptor and is required for Mic19 mitochondrial localization (11, 31) (Fig. 1A). Alternatively, H_2_S could affect protein stability and function via persulfidation, an oxidative cysteine modification, that results from the reaction of sulfide with an oxidized cysteine, e.g., a sulfenic acid or disulfide (16, 32).

**Figure 1.**
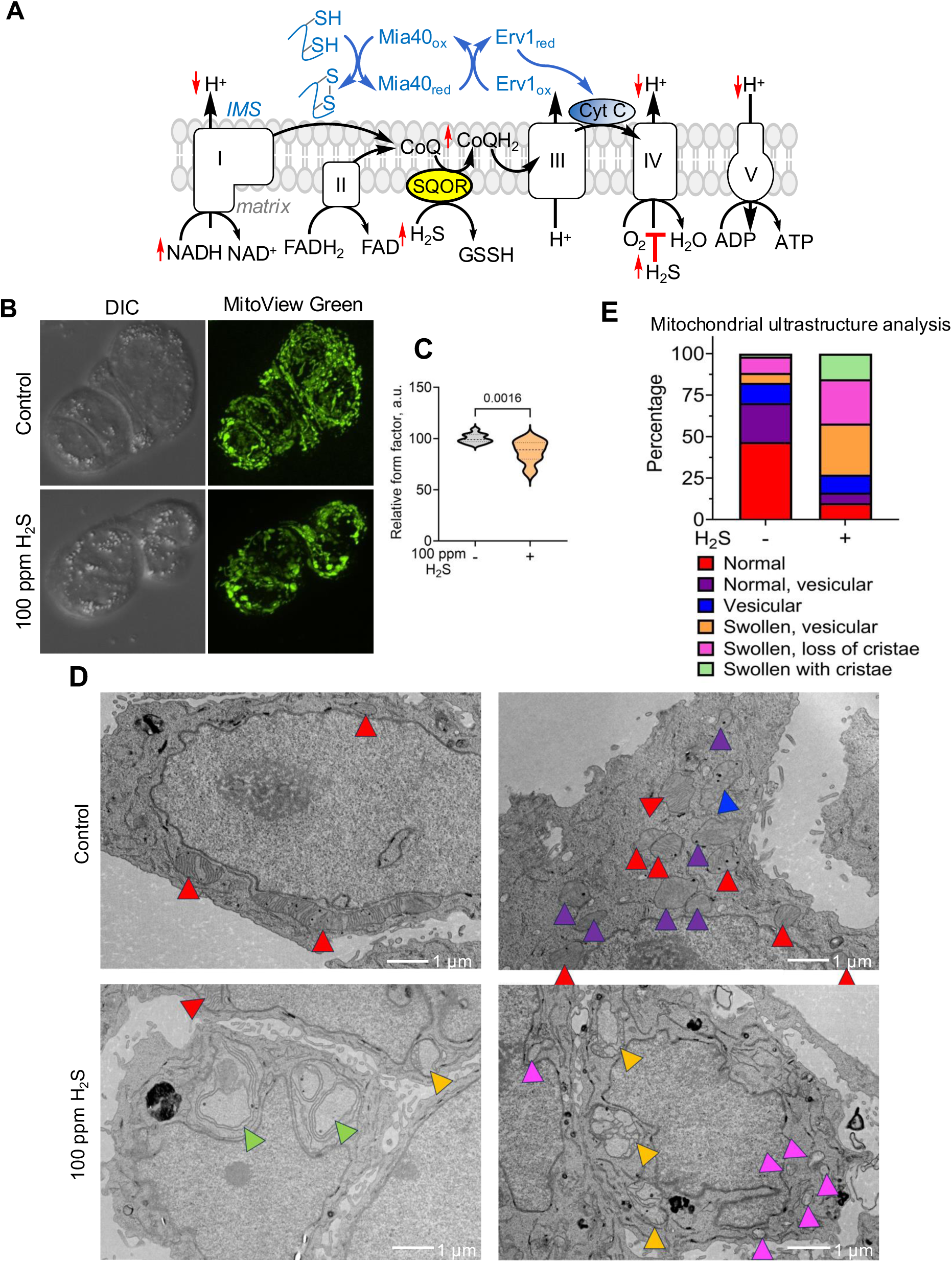
H_2_S induces cristae loss by disrupting MICOS. (**A**) Scheme illustrating the interactions of H_2_S with the ETC that lead to a reductive shift in cofactor pools and is predicted to do the same in the intermembrane space (IMS) impairing the Mia40-dependent pathway for oxidative protein folding. The red arrows indicate the anticipated direction of change in response to chronic H_2_S exposure. (**B**) Representative microscopy images of HT-29 cells stained with MitoView Green, reveals altered mitochondrial morphology and disrupted mitochondrial networking in response to H_2_S (100 ppm, 24 h). (**C**) Mitochondrial networks were estimated from the relative form factor of mitochondria in each control (n = 11) and sulfide (n = 13) image acquired from two independent experiments conducted in duplicate. **D**. Representative TEM images of HT-29 cells cultured ± H_2_S (100 ppm, 24 h) show changes in mitochondrial morphology that were characterized in 28 control and 46 H_2_S treated cells (scale bar: 1 µm). (**E**) The distribution of mitochondrial subtypes was expressed as a percentage of total mitochondria (n = 353 and 427 mitochondria without and with H_2_S treatment), using the same color scheme as in D.

Steady state tissue levels of H_2_S are low and vary between 10-80 nM (33, 34), but its concentration is significantly higher in colon lumen (0.2-2.4 mM) (35, 36), where it is influenced by diet and microbiota. Hereditary defects in the sulfide oxidation pathway enzymes lead to pathological elevation in sulfide and are associated with mitochondrial encephalomyopathies (37, 38). Exposure to a sublethal dose of sulfide (1000 ppm, 30 min) causes extensive mitochondrial damage in murine brain that is characterized by swelling and fragmentation, loss of cristae, and an increase in vesicular ultrastructure (39). The potential for physiologically relevant concentrations of sulfide to modulate mitochondrial form and function is however, unknown.

In the current study, we investigated the effects of chronic exposure to low sulfide (100 ppm, corresponding to 20 µM dissolved H_2_S in the culture medium) (40), on mitochondrial ultrastructure and networking. Under these conditions, cells exhibit S-phase cell cycle arrest and diminished mitochondrial networks, which was reversed upon removal of H_2_S from the atmosphere, signaling an adaptive response. H_2_S elicited membrane depolarization that was correlated with an increase in Oma1, a decrease in long Opa1 isoforms and a corresponding increase in short Opa1 isoforms. A closer examination revealed cristae loss and decreased CI and CIV activity associated with CI+CIII_2_+CIV_n_ and CIII_2_+CIV, which was correlated with a loss of the scaffold protein SCAF1 (or COX7A2L) from these supercomplexes. Finally, sulfide decreased levels of the ETC subunits GRIM19, NDUFB8, SCAF1 and COX7A2 and the CIV core subunits, MT-CO1 and MT-CO2. By providing a mechanistic framework linking sulfide to alterations in cristae dynamics, ETC stability and function, our study offers insights into how physiological fluctuations and pathological elevation in H_2_S might impact mitochondrial architecture and behavior.

## Results

### H_2_S remodels mitochondrial dynamics and cristae organization

Live cell imaging revealed profound effects of H_2_S on mitochondrial networking in HT-29 cells as revealed by form factor analysis, which is a measure of mitochondrial length and the degree of branching (41) (Fig. 1B,C). This effect was reversed within 48 h of sulfide removal (Fig. S1A). An H_2_S-induced decrease in mitochondrial networks was also seen in HEK293 (human embryonic kidney), HT1080 (human fibrosarcoma) and SW480 (human colon carcinoma) cells, which displayed more punctate mitochondria (Fig. S2-4). While these cell lines exhibited some variations in mitochondrial morphology in response to H_2_S, they exhibited an overall decrease in mitochondrial networking. For the remainder of the study, we focused on HT-29 cells for characterizing changes in mitochondrial ultrastructure in response to H_2_S.

Since mitochondrial dynamics are responsive to environmental and cellular stresses and central to functional regulation (42), we examined how sulfide exposure affects mitochondrial ultrastructure. Transmission electron microscopy (TEM) revealed mitochondrial swelling and cristae loss in sulfide- grown cells (Fig. 1D,E and Fig. S5). In contrast to untreated controls, in which most mitochondria (∼70%) exhibited normal or normal-vesicular morphology, >70% of sulfide-grown cells had swollen, swollen- vesicular mitochondria with partial or complete loss of cristae (Fig. 1E). In fact, only ∼10% of sulfide- grown cells retained normal morphology and >25% showed complete loss of cristae. Morphological changes were observed in most mitochondria irrespective of their size, and H_2_S-grown cells appeared to contain both smaller as well as much larger mitochondria relative to control cells.

Chronic H_2_S exposure (100 ppm, 24 h) diminished basal respiration in HT-29 cells by ∼90% and prevented subsequent sulfide-stimulated oxygen consumption, which requires a functional ETC (Fig. S6).

These effects were reversed within 48 h of the cells being moved to a growth chamber lacking sulfide as also observed at the level of mitochondrial morphology (Fig. S1A). We have previously shown that under these conditions, HT-29 cells exhibit decreased proliferation albeit, without a significant effect on cell viability (40). Cell cycle analysis revealed that sulfide (100 ppm, 24 h) increased the proportion of cells in the S-phase, which slowly redistributed over the next 48 h following removal of the sulfide from the growth chamber (Fig. S7). The G1-S transition is characterized by hyperfused mitochondria that are electrically continuous and have a higher ATP generating potential than at any other phase of the cell cycle (43). A combination of fragmented and hyperfused mitochondria was seen in sulfide grown cells reflecting S-phase enrichment.

### H_2_S disrupts inner membrane organization by destabilizing MICOS

MICOS stabilize cristae junctions, bridging the inner and outer membranes via interactions with SAM (sorting assembly machinery) in the outer membrane (Fig. 2A) (44). To further characterize sulfide-induced changes in cristae size and stability, expression of the MICOS subunits Mic19, Mic25, and Mic60 was assessed. Mic19 and Mic25 are essential for docking Mic60, the core assembly and maintenance component of MICOS, and loss of either protein leads to partial MICOS disassembly and irregular cristae morphology (45, 46). Chronic sulfide exposure decreased Mic19 and Mic60, but not Mic25 expression (Fig. 2B-D), which was correlated with an increase in isolated mitochondria as revealed by super resolution immunofluorescence imaging (Fig. 2E,F). Form factor analysis indicated disruption in mitochondrial networks, which was greater with the inner mitochondrial membrane marker, Mic60, than the outer membrane marker, Tom20 (Fig. 2G,H). These data suggest some selectivity of sulfide for targeting inner mitochondrial membrane proteins.

**Figure 2.**
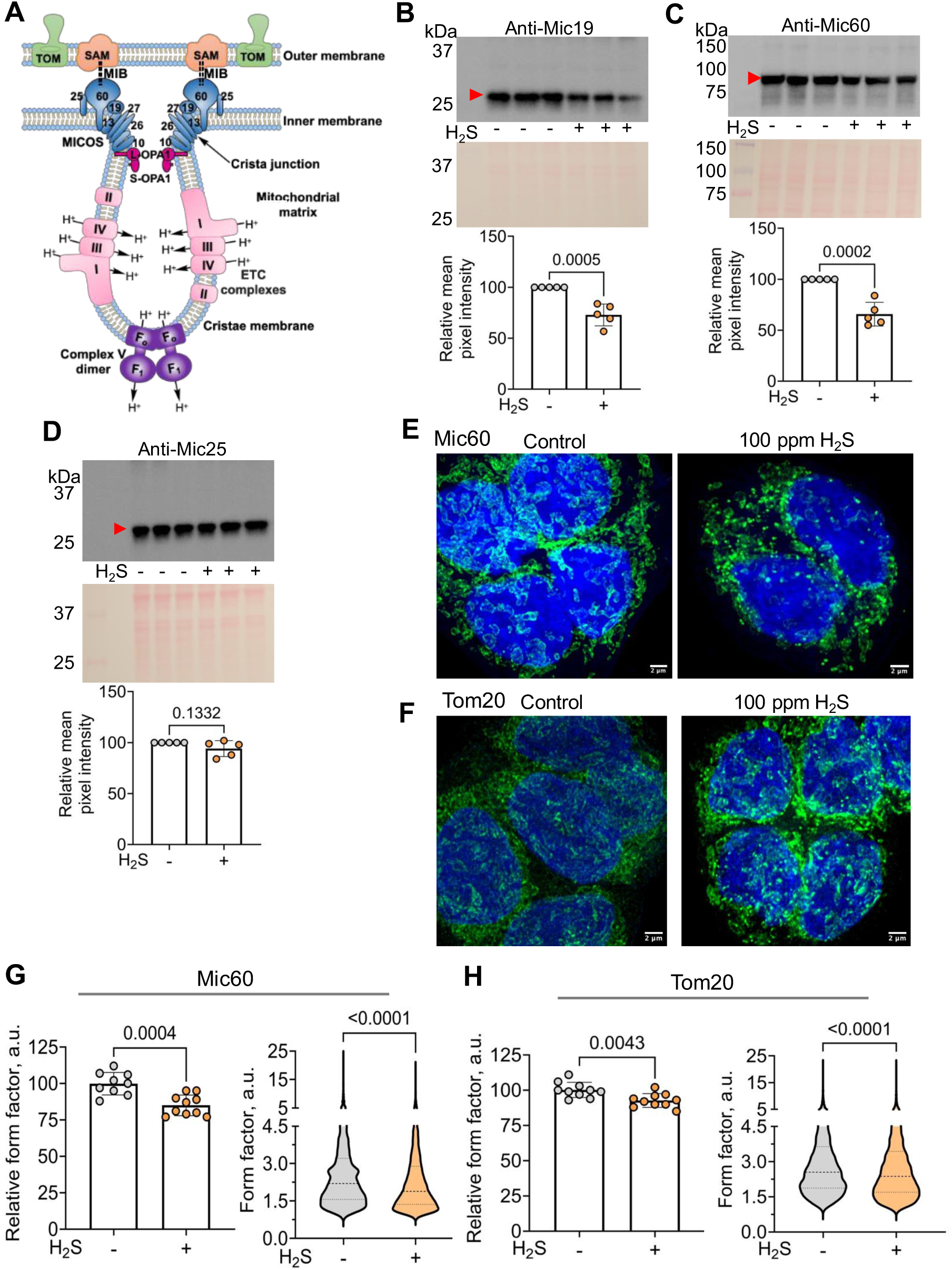
H_2_S induces cristae loss by disrupting MICOS. (**A**) Scheme showing organization of the mitochondrial membranes and cristae which was adapted from Ref (44). The MICOS complex subunits Mic10, Mic13, Mic19, Mic25, Mic26, Mic27, and Mic60 (denoted by numbers only) stabilize the cristae junction. MICOS links the inner and outer mitochondrial membranes through bridging interactions with the sorting and assembly machinery (SAM) complex, known as the mitochondrial intermembrane space bridging complex (MIB, dotted lines). The width of cristae junction is maintained by interactions between the long (L-) membrane bound, and short (S-) soluble isoforms of Opa-1. ETC complexes reside on the flanks and the F_1_F_o_ ATP synthase at the base of cristae, causing curvature. (**B-D**) Western blot analysis reveals that Mic19 and Mic60, but not Mic25, levels are significantly decreased in HT-29 cells exposed to H_2_S (100 ppm, 24 h). Ponceau red staining for equal loading (middle panels) and quantitative analysis of the blots (lower panels) are shown. Each symbol represents the average of 3 technical replicates from 5 independent experiments. (**E**,**F**) Representative immunofluorescence staining of Mic60 (E) and Tom20 (F) and DAPI in HT-29 cells cultured ± 100 ppm H_2_S for 24 h (scale bar: 2 µm). (**G,H**) Relative form factor per image (left panels) for Mic60 (n = 9, control; n = 10, H_2_S) (G) and Tom20 (n = 10, ± H_2_S) immunofluorescence staining, and the form factor (*right panels*) of each mitochondrion quantified in each image from two independent experiments conducted in duplicate. Two-sample unpaired t-test was performed for all statistical analyses.

### H_2_S influences inner membrane fusion by Oma1-dependent cleavage of Opa1

To elucidate the mechanism of sulfide-induced MICOS destabilization, we focused on the Oma1-Opa1 axis. Oma1 stabilizes MICOS (47) while the long and short isoforms of Opa1 regulate cristae junction width and inner membrane fusion (48). We hypothesized that ETC poisoning by H_2_S lowers mitochondrial membrane potential and upregulates and/or activates Oma1, which in turn, enhances proteolytic conversion of L- to S-Opa1 (9). Sulfide-grown HT-29 cells exhibited a pronounced decrease in mitochondrial membrane potential (Fig. 3A) and a small, by significant increase in Oma1 expression (Fig. 3B). Furthermore, while the L-Opa1 isoforms decreased, S-Opa1 isoforms increased, indicating enhanced proteolysis (Fig. 3C-E). Additionally, expression of Sirt4, the stress-responsive mitochondrial sirtuin that stabilizes L-Opa1 against proteolysis (49), was ∼40% lower in sulfide-grown HT-29 cells (Fig. 3F,G). These data are consistent with a model that increased Oma1 and decreased Sirt4 contribute to enhanced L-Opa1 cleavage in sulfide-grown cells. MEF (mouse embryonic fibroblast) cells in which Oma1 was knocked out, exhibited less intense mitochondrial staining with MitoView Green, which as in wild-type cells, increased in response to H_2_S (Fig. S8A). Form factor analysis revealed decreased mitochondrial networking in wild-type but not in Oma1 deleted cells in response to H_2_S (Fig. S8B).

**Figure 3.**
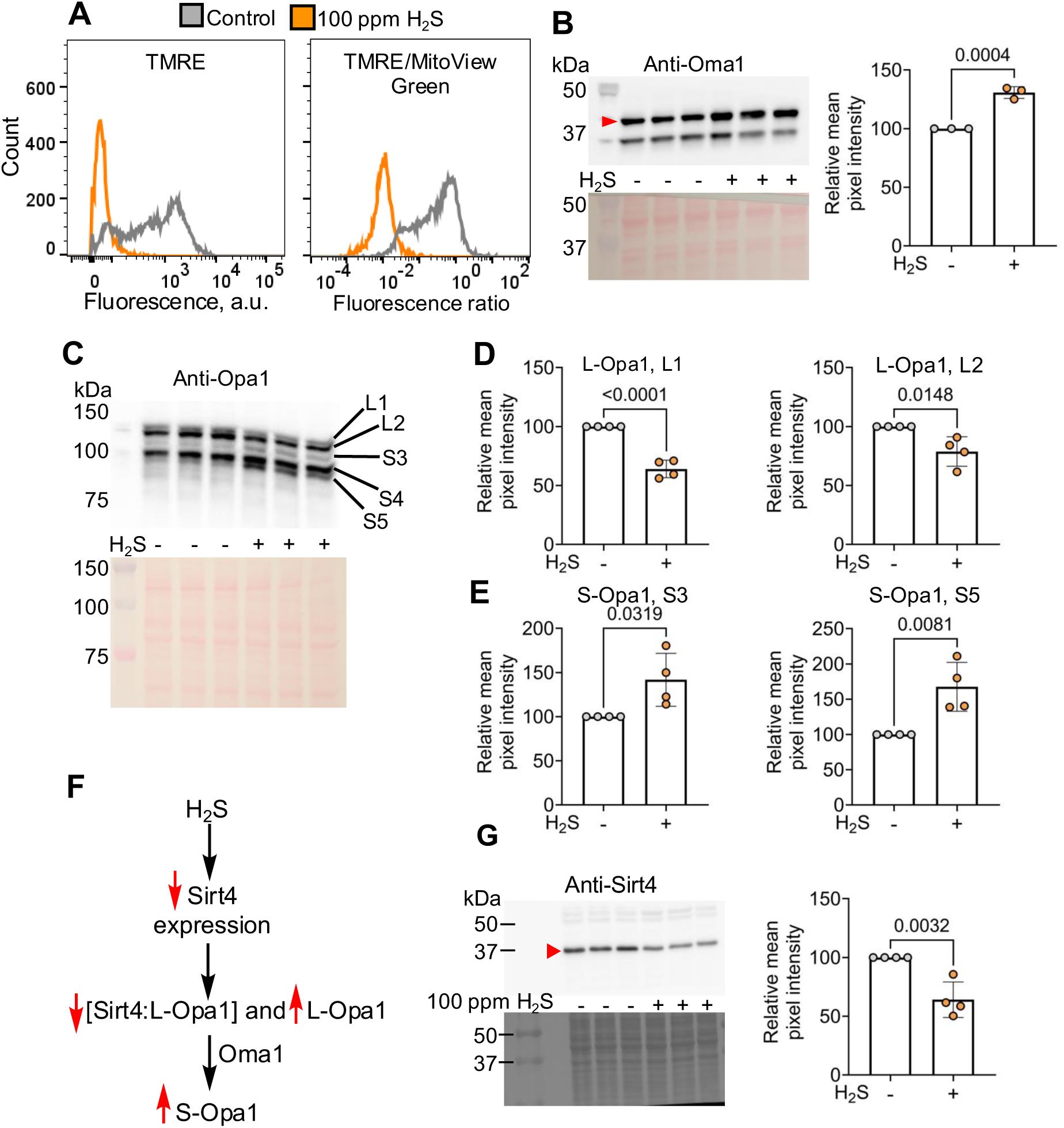
H_2_S lowers membrane potential and triggers OMA-dependent Opa1 cleavage. (**A**) Mitochondrial membrane potential in HT-29 cells cultured ±100 ppm H_2_S for 24 h was measured by TMRE staining (*left*) and normalized to mitochondrial mass estimated by MitoView Green staining (*right*). (**B**) Western blot data (*left*) and quantitative analysis (*right*) shows that H_2_S increases Oma1 expression (n= 3 independent experiments each in triplicate). (**C**) H_2_S leads to a redistribution of Opa1 isoforms (L an S denote long and short, respectively). (**D,E**) Quantitation of L1, L2 (D), S3, and S5 (E) Opa1 isoforms; (n= 4 independent experiments; each data point is the mean of 3 replicates). (**F**). Scheme illustrating the possible effect of H_2_S on Sirt4 on L-Opa1 stability. (**G**) Western blot analysis of Sirt4 levels (*left*) and quantitation (*right*) in HT-29 cells cultured ± 100 ppm H_2_S. (n= 3 independent experiments conducted in triplicate). A two-sample unpaired t test was performed for the statistical analysis and Ponceau staining for equal loading is shown below the respective western blots.

### H_2_S decreases mitochondrial respiration by disrupting supercomplex stability and assembly

We examined the effect of sulfide on supercomplexes since cristae morphology affects supercomplex assembly and efficiency (3, 50). Sulfide decreased CI and IV activity associated with high molecular weight respirasomes (Fig. 4A). A close-up revealed that CI activity was lost from the CS-respirasome (CI-CIII_2_-CIV_2_) but was partially retained in the A/C-respirasomes (CI-CIII_2_-CIV), which cannot be distinguished. A pronounced decrease in CIV activity in the Q-respirasome (CIII_2_-CIV) and the CIV dimer (CIV_2_) was also observed. Blue native PAGE (BN-PAGE) analysis of supercomplex abundance with markers for CI (NDUFV2), CIII (UQCRFS1), and CIV (COX4I1) revealed that CS- and Q-respirasomes and CIV_2_ were severely depleted in H_2_S-grown cells, while the A/C-respirasomes(CI-CIII_2_-CIV), were less impacted (Fig. 4B). In contrast, the CIII dimer (CIII_2_) and the CIV monomer (CIV) were markedly increased in intensity in sulfide grown cells. Each of the marker proteins was separately assessed by SDS-PAGE analysis of whole cell lysates to rule out that differences in their steady-state levels might have contributed to the analysis of supercomplex abundance (Fig. 4C-E).

**Figure 4.**
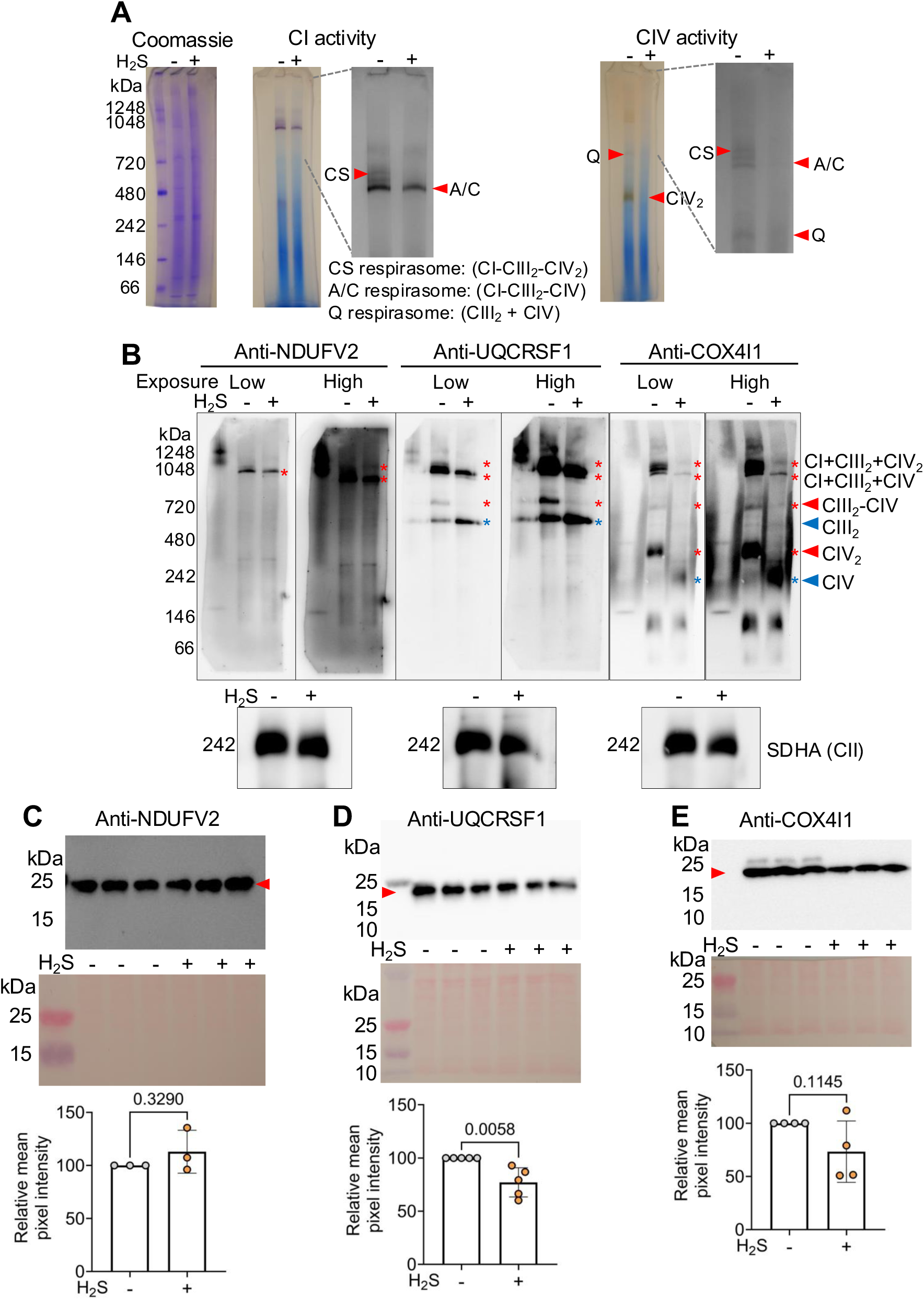
H_2_S disrupts respirasome and supercomplex stability. (**A**) Clear native PAGE of mitochondria from HT-29 cells (grown ± 100 ppm H_2_S, 24 h) stained for CI and CIV activity; Coomassie blue staining was used to demonstrate equal loading control (*left*). Close-ups of CI and CIV lanes (in gray scale) reveals lower activity for CS- and A/C-type respirasomes in H_2_S treated cells. (**B**) Whole cell lysate BN-PAGE and immunoblotting of NDUFV2 (CI), UQCRFS1 (CIII), and COX4I1 (CIV) (displayed at low and high exposure). The red and blue arrowheads indicate bands that are decreased or increased in abundance in response to H_2_S. Each blot was re-probed with SDHA to show equal loading (*lower)* and is representative of at least 3 independent experiments. (**C-E**) SDS-PAGE immunoblots of each protein probed in (B) shows that their levels were unchanged in H_2_S treated cells. Ponceau red staining (*middle*) for equal loading, and blot quantitation (*lower*) represent data from 3-5 independent experiments each conducted in triplicate. Two-sample unpaired t test was used for all statistical analyses.

Scaffold proteins are important for the interactions between CIII and IV in the CS-respirasomes (Fig. 5A) (51, 52). Western blot analysis revealed that the CIV scaffold proteins (SCAF1/COX7A2L and COX7A2), CI membrane anchoring proteins (GRIM19 and NDUFB8), as well as the CIV core subunits, MT-CO1 and MT-CO2, were decreased in H_2_S-grown cells (Fig. 5B-F). BN-PAGE immunoblot analysis indicated a significant loss of SCAF1/COX7A2L associated with the CS- and Q-respirasomes and loss of COX7A2 associated with the Q-respirasome and CIII_2_ (Fig. 5G). The latter assignments were confirmed with antibodies to CI (NDUFV2), CIII (UQCRSF1) and CIV (COX4I1). COX7A2 and NDUFB8, which are significantly lower in H_2_S-grown cells, are associated with the A- and C-respirasomes, while SCAF1/COX7A2L is not involved in their stabilization (Fig. S9).

**Figure 5.**
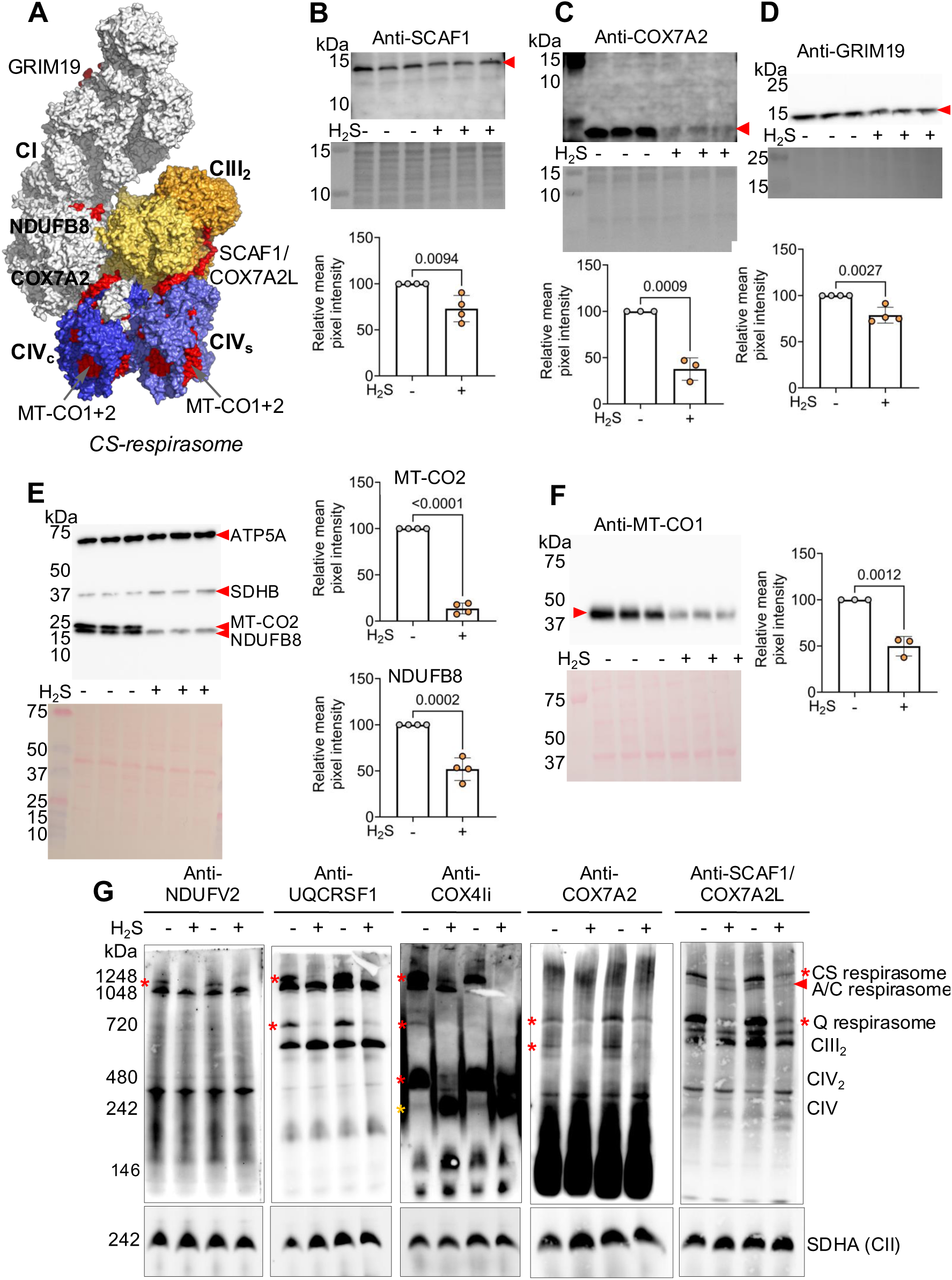
H_2_S leads to loss of complex I and complex IV scaffold protein from respirasomes. (**A**) Surface representation of the CS-respirasome (PDB 8PW7) depicting complexes I (gray), CIII_2_ (yellow) and CIV (blue), containing COX7A2 (CIV_c_) or SCAF1/COX7A2L (CIV_s_). The subunits that were probed in B-F and found to decrease in abundance in response to H_2_S are shown in red. (**B-F**) Western blot analysis of the complex IV scaffold proteins, COX7A2 (A) and SCAF1/COX7A2L (B), complex I inner membrane anchoring proteins GRIM19 (C) and NDUFB8 (D), and complex IV core subunits MT-CO2 (D) and MT-CO1 (E) show significant decreases in H_2_S-grown HT-29 cells (100 ppm, 24 h). Ponceau red staining for equal loading quantification are shown for each western blot. The data are from 3-4 independent experiments each conducted in triplicate. A two-sample unpaired t test was performed for all statistical analyses. (**G**) BN-PAGE immunoblots of HT-29 cell lysates (cultured ± 100 ppm H_2_S, 24 h) shows that SCAF1/COX7A2L (right panel) exhibits lower association with the CS-respirasome and with CIII_2_-CIV, while its levels in CIII_2_ is largely unchanged. COX7A2 is less abundant in CIII_2_ and the Q-respirasome. Anti-NDUFV2, UQCRFS1, and COXIV1i are shown side-by-side for verification of protein ID in the SCAF1/COX7A2L immunoblot. Red and yellow asterisks denote bands that are decreased or increased in response to H_2_S, respectively. The blots were re-probed for SDHA as an equal loading control (bottom panels). The data represent mitochondria isolated from 2 independent experiments.

## Discussion

Mitochondria are hubs for inter-organellar communication via physical contact sites with the endoplasmic reticulum, lysosomes, peroxisomes and lipid droplets (53), and mitochondrial plasticity is key component of cellular adaptation to stresses and changing metabolic needs (54). The mitochondrion has emerged as a prominent signaling hub for H_2_S, due to its twin potential to promote and inhibit ETC flux. In the gut, where host exposure to microbial-derived H_2_S is generally accepted to be high, the effective exposure of colonocytes, which are sheathed by a mucus layer, as well as the range of diurnal fluctuation in luminal H_2_S is not known. While insights into metabolic signaling, originating from the interaction of H_2_S with CIV, are beginning to emerge, little is known about how they are linked to mitochondrial ultrastructural changes. In this study, we demonstrate that chronic low-level H_2_S exposure is a significant modifier of mitochondrial ultrastructure and networking and delineate inner membrane protein targets of H_2_S. We posit that the observed effects originate with CIV inhibition by H_2_S, leading to membrane depolarization and a reductive shift in electron carrier pools. The downstream pleiotropic effects are mediated in part by promoting Oma1-dependent cleavage of Opa1, disrupting MICOS stability, and impairing supercomplex assembly. The impacts on mitochondrial networks and respiration are reversible under our experimental conditions, consistent with an adaptive response rather than progression along a cell death pathway.

TEM images revealed that H_2_S induced a significant increase in vesicular ultrastructure, a key indicator of cristae unfolding and mitochondrial swelling (Fig. 1D). A greater variability in mitochondrial size was observed in H_2_S-grown versus control cells (Fig. 1D and Fig. S5), which was correlated with a decrease in mitochondrial networking observed by live cell imaging of HEK293, HT-29, HT1080 and SW480 cells (Figs. S2-4). Levels of Mic60 and Mic19, two of three MICOS components examined in this study, were downregulated by H_2_S (Fig. 2B,C). Super resolution immunofluorescence imaging of Mic60 revealed a substantial decrease in form factor, a measure of mitochondrial networking (41), consistent with decreased inner membrane fusion (Fig. 1E,G).

Mic19 is a redox-regulated scaffold protein that associates with Mic60 in its disulfide form (11) and stabilizes the bridging interaction between MICOS in the inner, and Sam50 in the outer mitochondrial membranes (55). Mic19 knockdown increases mitochondrial fragmentation, lowers MT-CO2 expression, and increases vesicular ultrastructure (45), while Mic19 disulfide reduction leads to similar, albeit less pronounced changes (11). ETC inhibition by H_2_S leads to a reductive shift in redox cofactor pools (27, 28) and in cytochrome c (25), which are predicted to disrupt oxidative protein folding in the mitochondrial intermembrane space (Fig. 1A), impacting the stability of MICOS and other redox-sensitive proteins.

Mitochondrial flickering, i.e. pulsatile inner membrane depolarization, is regulated by the copper transport function of Slc25a3, a mitochondrial copper/phosphate carrier protein (56). Copper-dependent CIV (or cytochrome c oxidase) activity promotes flickering, which is attenuated by the inhibitor, sodium azide. We posit that H_2_S might be an endogenous modulator of mitochondrial flickering via CIV inhibition. We demonstrated that H_2_S induces membrane depolarization, which is a known trigger for Oma1 activation and consequent Opa1 proteolysis, a regulator of inner membrane fusion (48, 57) (Fig. 3A-E). Opa1 proteolysis could be further augmented by an H_2_S-dependent decrease in Sirt4, which helps stabilize L-Opa1 (49) (Fig. 3F-G). In cells lacking Oma1, sulfide failed to elicit a significant effect on mitochondrial networking (Fig. S8).

Structural changes in the mitochondrial inner membrane help orchestrate mitochondrial energy metabolism and fine tune supercomplex assembly (58). Defects in MICOS and a shift to short Opa1 isoforms have been shown to decrease supercomplex stability and decrease mitochondrial function (1), consistent with our findings that mitochondria in sulfide-grown cells exhibited diminished CI and IV activities associated with the CS- (CI-CIII_2_-CIV_2_) and Q- (CI-CIII2) respirasomes while the A/C- respirasomes were less affected (Fig. 4A). The CIV core subunits, MT-CO1 and MT-CO2, which were modestly decreased by acute H_2_S exposure (100 µM, 4 h) (26), were significantly diminished by chronic exposure (∼20 µM, 24 h). Additionally, depletion of CI (GRIM19, NDUFB8), CIII (SCAF1/COX7A2L) and CIV (COX7A2) subunits was observed in H_2_S-grown cells (Fig. 5B-F). COX7A2 is a component of CIV_c_, while SCAF1/COX7A2L is a component of CIV_s_ (51), and depletion of either is predicted to destabilize the CS-respirasome (Fig. 5A). This prediction was confirmed by loss of SCAF1/COX7A2L from supercomplexes in which CIV is a component but not from the CIII_2_ dimer (Fig. 5G). This conclusion is supported by depletion of the CS-respirasome and near complete loss of Q-respirasome (Fig. 4B), which require SCAF1/COX7A2L for stability (52). Sulfide-induced destabilization of MT-CO1 and MT-CO2 expression is consistent with disrupted mitochondrial bioenergetics and further highlights the link between loss of respiratory supercomplexes and impaired energy metabolism (Fig. S6). We posit that destabilization of supercomplexes could underlie the variable utilization of oxygen versus fumarate as a terminal electron acceptor by tissues (59, 60) and is governed by local differences in H_2_S as well as oxygen concentration.

In summary, our study reveals the potential for sulfide to modulate mitochondrial architecture and function, with possible metabolic implications for other organelles that communicate with this cellular energy hub. The observed changes in mitochondrial ultrastructure and network could be relevant in the pathology of diseases associated with elevated sulfide levels, e.g., inflammatory bowel disease (61, 62), as well as hereditary disorders that impair sulfide oxidation (37, 38). Our data suggest the possibility that at the low concentrations found in most cells, sulfide could be an endogenous regulator of mitochondrial dynamics possibly via inhibition of mitochondrial flickering-driven fusion.

## METHODS

### Materials

#### Cell lines

Human colorectal adenocarcinoma HT-29 and human embryonic kidney HEK293 cells were obtained from the American Type Culture Collection (ATCC). *Oma1*^-/-^ MEFs(12) and mito-RFP expressing human colorectal adenocarcinoma HT1080 and SW480 cells were generated as described previously (63).

#### Reagents

Na_2_S, nonahydrate (99.99% purity, (431648), protease inhibitor cocktail for mammalian tissue extract (P8340), dimethyl sulfoxide (D2650), reduced β-nicotinamide adenine dinucleotide (N8129), and cytochrome c from bovine heart (C2037-500mg) were from Sigma/Millipore. Dulbecco’s modified Eagle’s medium (DMEM) (with 4.5 g/L glucose, 584 mg/L glutamine, and 110 mg/L sodium pyruvate, (11995-065)), RPMI 1640 with glutamine (11875-093), RPMI + HEPES (22400105), phenol red free RPMI (11835030), fetal bovine serum (FBS, 10437-028), penicillin/streptomycin mixture (15140-122), 0.05% (w/v) trypsin-EDTA (25300-054), and PBS (10010-023) were from Gibco. KCl (7447-40-7) was from Acros Organics and Nonidet P40 substitute (74385) was from Fluka BioChemika. MitoView Green (70054) was purchased from Biotium. Methanol (A452-4), Tween20 (BP337500), nitro blue tetrazolium chloride (N6495), tetramethylrhodamine ethyl ester perchlorate (TMRE) (T669), and NativePAGE^TM^ 3-12% gels (10-well, BN1001BOX; 15-well, BN1003BOX), and the NativePAGE™ Sample Prep Kit (BN2008), containing NativePAGE™ 4x Sample Buffer or pinky lysis buffer (BN2003) and NativePAGE™ 5% G-250 Sample Additive (BN2004) were purchased from Fisher. EGTA (33963) was from Cayman Chemical. 10% Pre-cast tris-glycine gels (4561033), PVDF membranes (162-0177), thick blot filter paper (1703932), Clarity ECL and Clarity Max ECL substrates (1705061 and 1705062, respectively) were purchased from Bio-Rad. Polystyrene round-bottom tubes (5 mL) with cell-strainer caps used for FACS analysis was from BD Biosciences (720035). High glass bottom dishes (35 mm, 81158) for microscopic imaging of live cells were purchased from Ibidi. A propidium iodide flow cytometry kit (ab139418) for DNA staining and FACS analysis was purchased from Abcam. Cylinders with H_2_S in N_2_ (500 L, “A33” 5000 ppm H_2_S with 1% accuracy; #3130) or breathing air containing 5% CO_2_ were from Cryogenic Gases (Detroit, MI, USA).

Reagents for TEM sample preparation were purchased were from Electron Microscopy Sciences, including 3% glutaraldehyde and 3% paraformaldehyde in 0.1 M cacodylate buffer (15950), 0.1M cacodylate buffer (11654), 1.5% potassium ferrocyanide in 0.1 M cacodylate buffer (19150), osmium tetroxide in cacodylate buffer (25154-10), 0.1 M acetate buffer (11482-42), 2% uranyl acetate in 0.1 M acetate buffer (22400-4), 200 proof ethanol (15055), HPLC grade acetone (022928.K2), and Spurr’s Resin (14300).

#### Antibodies

Anti-Grim19 (1:1000; ab110240), anti-MT-CO1 (1:1000; ab14705), anti-MT-CO2 (1:1000; ab110258), anti-NDUFV2 (1:1000; AB183715), anti-OxPhos (1:1000; ab110411), and anti-UQCRFS1 (1:1000; ab14746) were from Abcam. Anti-COX4I1 (1:1000; VPA00544) was purchased from Bio-Rad, and anti-Opa1 (1:1000; 80471) was purchased from Cell Signaling Technology. Anti-COX7A2 (1:200; 18122-1-AP), anti-COX7A2L/SCAF1 (1:500; 11416-1-AP), anti-Mic19/CHCHD3 (1:5000; 25625-1-AP), anti-Mic25/CHCHD6 (1:2000; 20639-1-AP), anti-Mic60/Mitofilin (1:5000; 10179-1-AP), and anti- Sirt4 (1:1000; 66543-1-IG) were purchased from ProteinTech. Anti-Oma1 (1:1000; SC515788) was purchased from Santa Cruz Biotechnology. Anti-Tom20 (1:1000; MABT160) was purchased from Sigma/Millipore. Anti-rabbit horseradish peroxidase-linked IgG and anti-mouse IgG, horseradish peroxidase linked antibodies (1:10,000; NA944V and NA931, respectively) were from GE Healthcare.

#### Cell culture

HT-29 cells were maintained in RPMI 1640 medium. HEK293, HT1080, SW480, and MEF cells were maintained in DMEM medium. Both RPMI and DMEM media were supplemented with 10% FBS along with 100 units/mL penicillin and 100 µg/mL streptomycin. All cells were maintained at 37°C with ambient O_2_ and 5% CO_2_. During culture in the sulfide growth chamber, cells were maintained in the same culture medium but with twice the volume of medium to prevent acidification as described (40).

#### Transmission electron microscopy

Replicate 10 cm plates were seeded with 1x10^7^ HT-29 cells and cultured overnight. The following day, fresh 20 mL medium per plate was added, and cells were cultured for 24 h ± 100 ppm H_2_S. Cells were fixed in 3% glutaraldehyde and 3% paraformaldehyde in 0.1 M cacodylate buffer (CB, pH 7.2) overnight at 4°C. Then, cells were washed 3 x 15 min with CB, and subjected to osmification with 1.5% K_4_Fe(CN)_6_ + 2% OsO_4_ in 0.1 M CB for 1 h at room temperature, followed by 3 x 5 min washes with 0.1 M CB buffer at room temperature. The cells from replicate plates were scraped, combined, and centrifuged at 300 rpm for 10 min. Prewarmed 4% agarose was added to the cells pellets and cooled on ice. Next, the samples were washed 3 x 5 min with 0.1 M sodium acetate buffer (pH 5.2) at room temperature, then stained for 1 h stain with 2% uranyl acetate in 0.1 M sodium acetate buffer (pH 5.2). After 2 x 5 min additional 0.1 M sodium acetate buffer washes and a 1 x 5 min distilled water wash, the samples were dehydrated by serial 15 min washes in ethanol (30, 50, 70, 80, 90, 95, and 100%) before washing in acetone and infiltrating with Spurr’s resin with 1, 2, and 16 h room temperature treatments with 2:1, 1:1 and 1:2 acetone:Spurr’s Resin mixtures, respectively. Next, the samples were placed in 100% Spurr’s resin for 24 h at room temperature before embedding in the resin to polymerize at 70°C for 24 h. The resulting blocks were then sectioned at 70 nm thickness using a Leica EM UC7 ultramicrotome and imaged at 60 kV using a JEOL 1400+ transmission electron microscope. Mitochondrial ultrastructure classification was done by blinding all images, and then manually annotating with the following classifications as described previously (64): normal, normal-vesicular, vesicular, swollen-vesicular, swollen, and swollen with cristae. Mitochondria (353) were analyzed from 28 control cells, and 427 mitochondria were analyzed from 46 H_2_S cultured cells.

### SDS-PAGE Western blot analysis

#### Sample preparation

Cells were seeded at 2x10^6^ cells per 6 cm plate and cultured overnight before replacing with 8 mL fresh cell culture medium and culturing for 24 h ± 100 ppm H_2_S in the sulfide growth chamber as described (40). To harvest, cells were washed once with 5 mL PBS followed by addition of 0.5 mL 0.05% trypsin for 5 min at 37°C before collection in 1 mL RPMI medium. Cell suspensions were centrifuged at 1600 x *g* for 5 min, the pellet was washed with 1 x 1 mL ice cold PBS, resuspended in 300 µL non-denaturing lysis buffer with protease inhibitor (0.5% (v/v), Nonidet P40 substitute, 25 mM KCl, 20 mM HEPES, pH 7.4), and then frozen until use.

#### Development of SDS-PAGE western blots

Frozen cell pellets were lysed by three freeze-thaw cycles and centrifuged at 13,000 x *g* for 10 min. Protein content in the supernatant was determined using the Bradford reagent (Bio-Rad). Cell lysates (10-30 µg) were electrophoresed using precast Bio-Rad 10% tris-glycine SDS gels, transferred to PVDF membranes, blocked for 1 h with 5% milk in Tris buffered saline with 0.3% Tween 20 (TBST). Membranes were soaked overnight in TBST, 5% (w/v) milk containing diluted antibodies and then quickly washed twice with TBST followed by 4 x10 min washes with TBST. Membranes were exposed for 90 min to the secondary antibody (horseradish peroxidase-linked anti-rabbit or anti-mouse IgG used at a 1:10,000 dilution in TBST, 5% milk). The membranes were quickly washed twice with TBST followed by 5 x 10 min TBST washes and two rinses with TBS before treating with clarity ECL substrate (Bio-Rad). Signals were detected using a Bio-Rad ChemiDoc Imaging System. All images were exported as 16-bit TIF files and imported into Fiji (65) for semi-quantitative analysis, which was performed by drawing equal sized rectangles over each band to estimate its mean pixel intensity. The background was subtracted from each band.

### Native gel electrophoresis of mitochondrial proteins

#### Crude mitochondrial isolations

P2 crude mitochondrial fractions were isolated as previously described with modifications (66). Cells were seeded at 5-6 x10^6^ cells per 10 cm plate in triplicate and cultured overnight before replacing with fresh 20 mL medium and culturing for 24 h ± 100 ppm H_2_S. Cells were harvested by washing each plate once with 8 mL PBS, treating with 1 mL 0.05% trypsin for 5 min at 37 °C before harvesting in 6 mL RPMI medium per plate. Cells were pooled in 50 mL conical tubes, pelleted for 5 min at 1600 x g at 4°C, washed once with 25 mL ice-cold PBS, and pelleted again for 5 min at 1600 x g at 4°C. The cell pellet was frozen for later use or resuspended in 3 mL ice-cold CP-1 buffer (100 mM KCl, 50 mM Tris, 2 mM EGTA, pH 7.4 + protease inhibitor) for homogenization on ice by 20 passes with a glass homogenizer with a teflon tip, followed by 5 passes through a 27-gauge needle using a 5 mL syringe. The homogenate was pelleted at 700 x g for 10 min at 4°C. The top 90% of the mitochondria-containing supernatant was collected while the pellet was suspended in an additional 3 mL of CP-1 buffer and re-homogenized by 20 additional passes with the glass homogenizer and 5 additional passes through the 27-gauge needle. The resulting homogenate was pelleted at 700 x g for 10 min at 4°C, and the top 90% of the second supernatant batch was pooled with the first and aliquoted in 6x1.7 mL conical tubes and pelleted at 10,400 x g for 10 min at 4°C. The resulting mitochondria-enriched P1 pellets were consolidated in 3x1.7 mL conical tubes using 3 mL of CP-1 buffer and centrifuged again at 10,400 x g for 10 min at 4°C to obtain crude P2 pellets. The latter were combined into one slurry using 450 µL CP-1 buffer, which was pipetted up and down with a 200 µL pipet before using 5 µL of the mitochondrial slurry to determine protein content, using the Bradford reagent (Bio-Rad) and a calibration curve generated with 0.5-4 mg/mL BSA standards in CP-1 buffer. The mitochondria were then aliquoted in 100 µg fractions for later use and stored at -80°C.

Before use, 100 µg aliquots of P2 mitochondria were reisolated by thawing on ice and then centrifuged at 12,600 x g for 5 min. The supernatant was removed and the pellet was resuspended in 54 µL 1x pink lysis buffer (Fisher) + Sigma protease inhibitor cocktail. Next, 12 µL of 5% digitonin (Sigma) was added to achieve a ratio of protein/detergent of 1:6, the samples were mixed by pipetting, and incubated on ice for 15 min before centrifugation for 20 min at 20,000 x g. The top 60 µL of supernatant, which contained solubilized mitochondrial proteins, was transferred into a pre-labeled tube. Next, 4 µL of blue native loading buffer (Invitrogen, BN20041) was added, yielding a final concentration of 1.4 µg/µL in each sample for loading in precast Bis-Tris 3-12% gradient gels (Invitrogen, BN2011B × 10) for either clear native (CN) or blue native (BN) PAGE.

#### Preparation of whole cell lysates

Digitonin permeabilized cell lysates for BN-PAGE analysis were prepared as previously reported with slight modifications (67). Cells were seeded at 5-6 x10^6^ cells per 10 cm plate in quadruplicate and cultured overnight before replacing with 20 mL fresh culture medium and placing them a growth chamber for 24 h ± 100 ppm H_2_S. Cells were harvested by washing each plate with 8 mL PBS before treating with 1 mL 0.05% trypsin for 5 min at 37 °C. The trypsinized cells were collected in 7 mL RPMI medium/plate and centrifuged at 1600 x g for 5 min. The cell pellet was suspended in 8 mL ice-cold PBS and a 100 µL aliquot was diluted 1:1 (v/v) with trypan blue for cell counting. An aliquot (20 µL) was used for counting in a Cellometer (Nexelcom). The remaining cell suspension was centrifuged at 1600 x g for 5 min in several 1.7 mL conical tubes to obtain 2.5 x 10^6^ cell aliquots; each was washed once with 1 mL ice cold PBS, resuspended in 200 µL ice-cold PBS, and then spiked with 7.5 µL of 5% digitonin to yield a final concentration of 1.8 mg/mL digitonin. Cells were mixed by pipetting and incubated on ice for 10 min before adding 1 mL ice-cold PBS and spinning for 5 min at 20,000 x g. The supernatant was aspirated, and cells were washed with 1 mL ice-cold PBS before centrifuging for 5 min at 20,000 x g. The resulting cells pellets, which are permeabilized cells, were frozen at -80 °C until later use.

Before use, three aliquots of isolated, permeabilized whole cell lysates were thawed on ice for 30 min, consolidated in 310 µL of pink lysis buffer + protease inhibitor. The protein content of the samples was measured by the Bradford assay, and ∼310 µg protein was spiked with 25 µL of 5% digitonin (w/v) to a final protein/detergent ratio of 1:4. The samples were mixed by pipetting and then incubated on ice for 10 min before centrifuging for 30 min at 20,000 x g. The top 270 µL of the supernatant was transferred to pre-labelled tubes containing 20 µL of NativePAGE™ 5% G-250 Sample Additive (Fisher, BN2004). The final concentration of solubilized protein for BN-PAGE was ∼0.9 µg/µL.

#### Development of BN-PAGE western blots

Solubilized mitochondrial proteins were separated by loading 10 µL P2 crude mitochondrial sample (14 µg per lane) or 40 µL of whole cell lysate sample (36 µg per lane) onto precast Bis-Tris 3-12% gradient gels (Invitrogen, BN2011B × 10) in the NativePAGE™Novex Bis-Tris Gel System according to manufacturer’s recommendations for organelle protocols and following key considerations as described previously (68). Proteins were separated with NativePAGE™ anode buffer (Invitrogen, BN2001) and dark blue cathode buffer (Invitrogen, BN2002) for 1 hat 150 V at 4 °C before switching to light cathode buffer (Invitrogen, BN2002) to run overnight at 20-30 V at 4°C. Proteins from the resulting gels were transferred to PVDF membranes (Bio-Rad) using the wet transfer technique with the XCell II™ Blot Module overnight at 4°C. After transfer, the membranes were fixed for 5 min in 8% acetic acid, washed twice with water for 5 min, and then clipped to air dry. Next, each blot was rinsed with methanol (3 x 5 min) and water (3 x 5 min), and blocked in 5% milk in TBST for 30 min before blotting with the indicated primary antibodies according to the manufacturers’ recommendations. Secondary anti-mouse HRP antibodies were the same as listed for the SDS-PAGE analysis, but BioRad Clarity Max (1705062) was used as the ECL substrate to visualize the native PAGE immunoblotted bands.

### CN-PAGE in-gel activity assays

The in gel activity assays were performed at previously described (68), by loading 20 µg (1.4 µg/µL) of solubilized P2 crude mitochondria in 10-well, 3-12% Bis-Tris gradient gels in the NativePAGE™ Novex Gel system described above, but by first separating with NativePAGE™ anode buffer and light blue cathode buffer for 30 min at 150 V before switching to clear cathode buffer to avoid excessive blue color of the Coomassie dye on the gel that would interfere with the color of the activity. The gel was run for additional 2.5-3 h at 250 V for better separation of the SC bands (68). Complex I activity was stained with 0.1 mg/mL NADH (2 mg) and 2.5 mg/mL nitrotetrazolium blue chloride (50 mg) in 20 mL of 2 mM Tris-HCl, pH 7.4 for 20-30 minutes before stopping the reaction with 10% acetic acid, rinsing with water, and then imaging (68). Complex IV activity was stained with 2mg/mL diaminobenzidine, 4 mg/mL bovine cytochrome c in 45 mM phosphate buffer, pH 7.4 for an hour before stopping the reaction with 10% acetic acid, rinsing the gel with water, and imaging (68).

### Mitochondrial networking analysis

*MitoView Green stained cells.* HT29, HEK293, MEF, and MitoTag-RRP expressing HT1080 and SW480 cells were plated at 50,000-100,000 cells per 35 mm high glass bottom dishes (Ibidi, 81158) and cultured overnight before replacing the medium and placing in the growth chamber for 24 h ± 100 ppm H_2_S. To stain HT29, HEK293, and MEF cells, the culture medium was replaced with serum-free RPMI containing 50 nM MitoView green added from a 200 µM stock in DMSO. The cells were stained in the growth chamber for 30 min before washing the cells twice with serum free RPMI and replacing the medium with phenol red free RPMI + FBS with penstrep and cultured for an additional 60-90 min before imaging. The MitoTag-RRP expressing cells had been previously validated to have mitochondrial localization of RFP and were not stained (63). Cells were imaged on Nikon X1 Yokogawa Spinning Disk Confocal microscope with CO_2_ and temperature regulation, 63x oil objective with a 1.49 numerical aperture, equipped with an Andor DU-888 monochrome camera for widefield imaging and a 488 nm laser for exciting the MitoView Green dye or a 561 nm laser for exciting mitoRFP.

#### Image analysis and form factor calculations

The form factor (a function of both mitochondrial length and branching, equal to perimeter2/4π × area) was assessed to measure mitochondrial networking using FIJI as described previously (69). For this, maximal intensity projections of the fluorescence intensity images were made, and equal intensity thresholding was used for all analyzed images. The threshold images were used to identify mitochondria, whose area and perimeter were exported into an excel file to calculate the form factor for each mitochondrion. The mitochondrial data were exported in bulk to quantify the form factor of every identified mitochondria, or each image’s data averaged per image to quantify the average form factor per image. The data were used to generate violin plots in Prism.

### Immunofluorescence microscopy

Cells (25,000-50,000 cells/well in a 24-well plate) were seeded in pre-sterilized (190 proof ethanol, 10 min), poly-d-lysine precoated (1 h, 37 °C) coverslips and cultured for 24 h. The next day, the medium was changed, and cells were cultured for 24 h ± 100 ppm H_2_S before fixation with 4% paraformaldehyde for 15 min. Cells were washed with PBS, permeabilized with 0.25% Triton X-100 in PBS for 5 min, blocked in CST blocking buffer (5% Donkey/Goat serum/BSA, 0.3% Triton X-100 in PBS) for 30 min, and then incubated for 1 h with primary antibodies against Mic60 or Tom20 diluted 1:200 in buffer (1% BSA, 0.3% Triton X-100 in PBS). The samples were washed 3 x 5 min with 1% BSA, 0.3% Triton X-100 in PBS before treating with secondary 488 Donkey/Goat-anti-rabbit antibody (Invitrogen) diluted to 1:500 in buffer (1% BSA, 0.3% Triton X-100 in PBS) for an hour at room temperature in a humidified chamber. The samples were washed 3 x 5 min with 1% BSA, 0.3% Triton X-100 in PBS before letting the coverslips dry before mounting them on glass slides using ProLong Gold + DAPI (Fisher) and imaged using a Zeiss LSM 980 Airyscan 2 microscope and detector equipped with a 63x oil objective with a 1.4 numerical aperture. Post-processing was done with Zen 3.4 (Blue edition), and maximum intensity projections and segmenting of mitochondria for form factor analysis was done in FIJI.

### Flow cytometry analysis

FACS analysis as conducted using the Bio-Rad Ze5 multi-laser, high speed cell analyzer operated with the Everest software package at the University of Michigan Flow Cytometry Core Facility. All data were analyzed using FlowJO (v10.8.1).

#### TMRE membrane potential measurements

HT-29 cells were seeded at 500,000 cells per well in 6- well plates and grown overnight before replacing the medium and placing cells in the growth chamber ± 100 ppm H_2_S for 24 h. The medium was replaced with serum free RPMI containing 50 nM TMRE and 50 nM MitoView Green (in DMSO) and staining was continued for 30 min ± 100 ppm H_2_S culture, before washing the cells twice with room temperature RPMI, and replacing the medium with phenol red free RPMI + FBS with penstrep. The cells were cultured in this medium for an additional hour before harvesting to minimize nonspecific binding of the dyes. Then, the medium was aspirated, and cells were collected by scraping in 500 µL ice-cold PBS, suspended by pipetting, filtered through 5-mL BD round bottom falcon tubes with cell-strainer caps, for FACS analysis. FlowJO (v10.8.1) was used to conduct gating and ratio TMRE:MitoView Green fluorescence for each analyzed cell, and to plot histograms of the raw and ratioed TMRE fluorescence.

#### Cell cycle analysis

To evaluate the reversibility of H_2_S-induced changes in the cell cycle, 2 x10^6^ HT- 29 cells per 6 cm plate (in duplicate) were cultured overnight. The next day, fresh 8 mL medium was added to each plate before placing them in the growth chamber ± 100 ppm H_2_S for 24 h. Then, cells were either harvested or allowed to recover in the absence of H_2_S for 24-48 h with fresh medium changes every 24 h. The conditioned medium was collected in 15 mL conical tubes and cells were washed with PBS, which was collected in the same tube. Next, the cells were treated with 0.5 mL 0.05% trypsin for 5 min at 37°C and harvested with 1 mL RPMI medium, pooled with the conditioned medium and PBS wash, and centrifuged at 500 x g for 5 min at 4°C. Cells were washed once with 1 mL ice-cold PBS by gentle suspension and subsequent cell pelleting at 500 x g for 5 min at 4°C. Cells were then fixed in 66% ethanol on ice by suspending the cell pellet in 400 µL of ice cold PBS and slowly adding 800 µL of ice cold absolute ethanol, which was mixed by pipetting. The fixed cells were stored at 4°C until all time points were harvested before proceeding with DNA staining and FACS analysis with the propidium iodide flow cytometry kit for cell cycle analysis (ab139418 from Abcam), following the manufacturer’s protocols.

### Oxygen consumption rate (OCR) measurements

All OCR measurements were performed on the Oroboros Instruments Corp respirometer at 37 °C with a stirring rate of 750 rpm. Cells were harvested from 10 cm plates, washed with 1 x 8 mL PBS prior to digestion with 1 mL trypsin (0.05%) at 37°C for 5 min. Cells were collected in 7 mL of the cell culture medium and centrifuged at 1600 x *g* for 5 min, and pellets were resuspended in 1 mL modified PBS (MPBS) or DPBS + 5 mM glucose + 20 mM HEPES, pH 7.4. Cell suspensions were transferred to pre-weighed tubes, centrifuged at 1600 x *g* for 3 min and the supernatant was carefully aspirated with a 2 µL tip fixed to a vacuum line. The wet weight of the pellet was determined to prepare 5 % (w/v) cell suspensions in MPBS, which were kept on ice and used for dilution to 1% (w/v) suspensions in MPBS for OCR experiments.

Analysis of OCR traces was performed using DatLab v6 (Oroboros Instruments, Austria) and replotted in Origin 7.0. Recovery time following Na_2_S injection is defined as the time taken by cells to return to a new stationary basal OCR.

### Statistics

Statistical analysis for pairwise comparisons was performed using the two-sample unpaired t-test.

## ACKNOWLEDGEMENTS

This work was supported in part by grants from the National Institutes of Health (F32GM140694 to DH, GM130183 to RB, and GM131701-01 to OK).

## DATA AND MATERIALS AVAILABILITY

All data are available in the manuscript or supplementary materials.

## AUTHOR CONTRIBUTIONS

D.A.H. and R.B. designed research; D.A.H. performed the research; B. Chen assisted with microscopy. O.K., B. Chen, and Y.S. contributed new reagents, D.A.H., B. Cunniff, and R.B. analyzed data; D.A.H. and R.B. wrote the manuscript while all authors reviewed and edited it.

---

**Figure S1.**
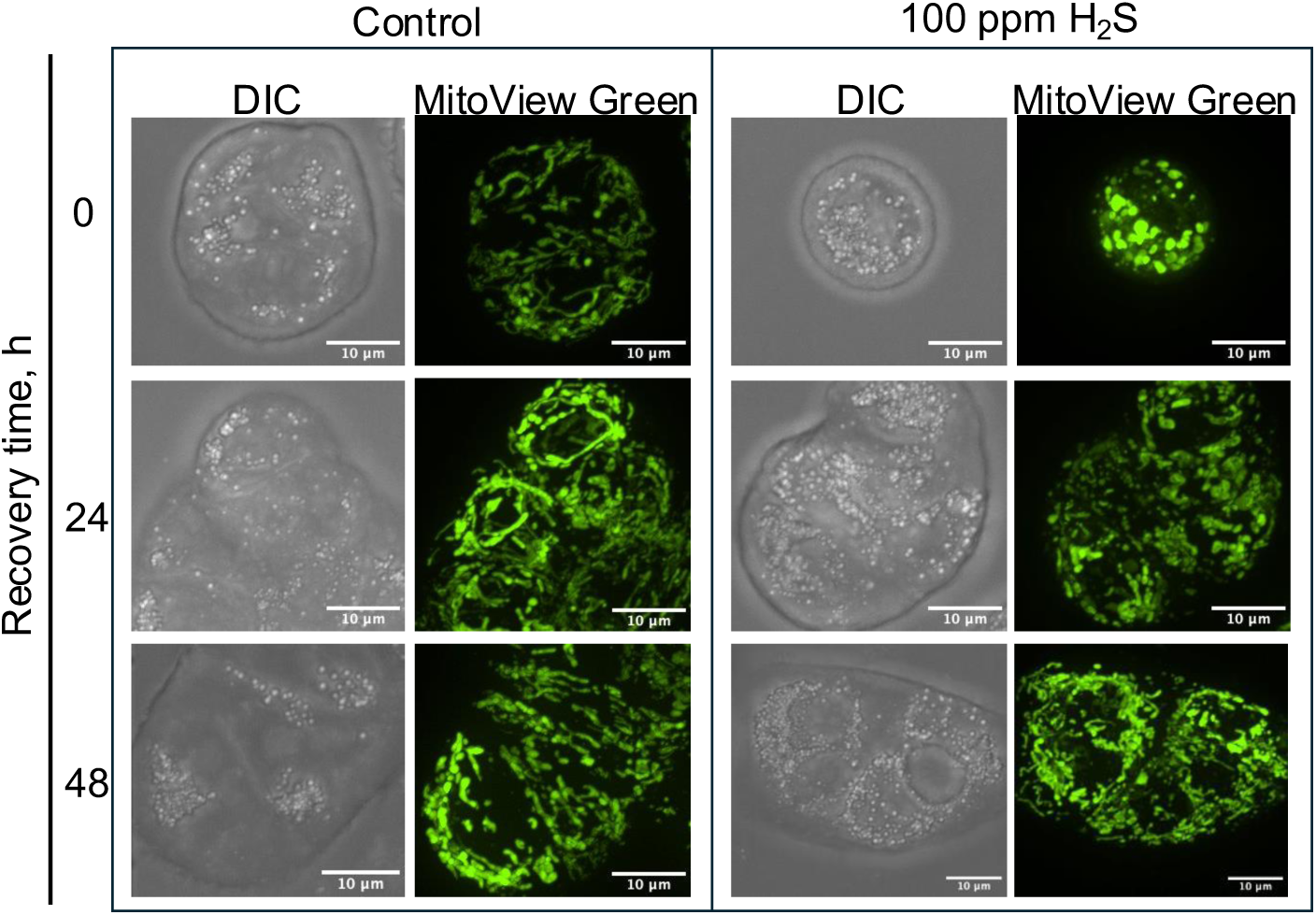
H_2_S induced mitochondrial swelling is reversible. Mitochondrial morphology is altered by H_2_S (100 ppm, 24 h) but recovers by 48 h after return to growth in the absence of H_2_S under normal growth conditions. Mitochondria were stained with MitoView. And data are representative of 2 independent experiments.

**Figure S2.**
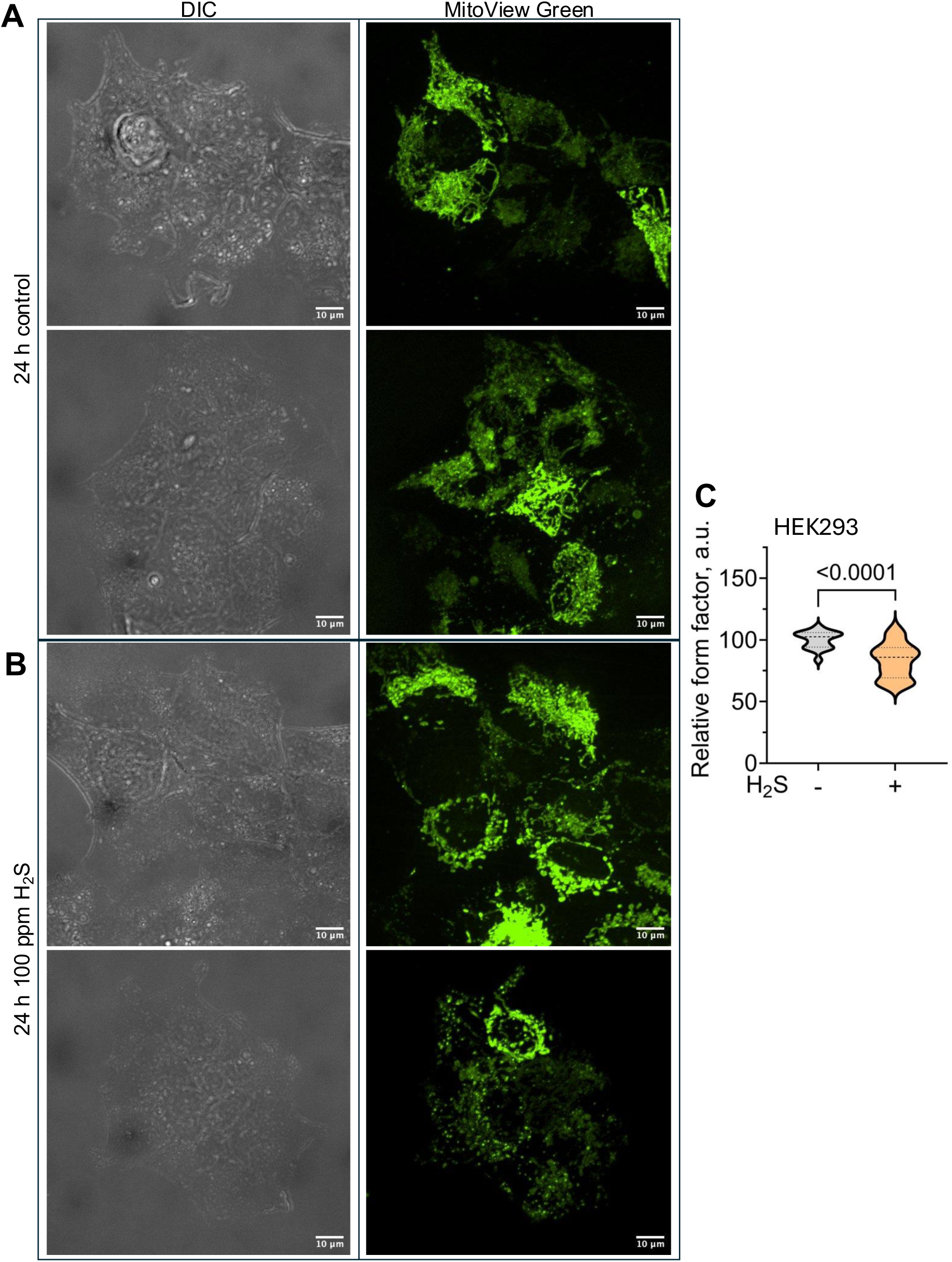
Chronic H_2_S decreases mitochondrial networking in HEK293 cells. (**A,B**) Mitochondrial morphology in control (A) versus (B) H_2_S-grown cells (100 ppm, 24 h) altered mitochondrial morphology and decreased mitochondrial networking. Two representative images are shown to illustrate variations in response to H_2_S. (**C**) Form factor analysis revealed decreased mitochondrial networking (n >20 images).

**Figure S3.**
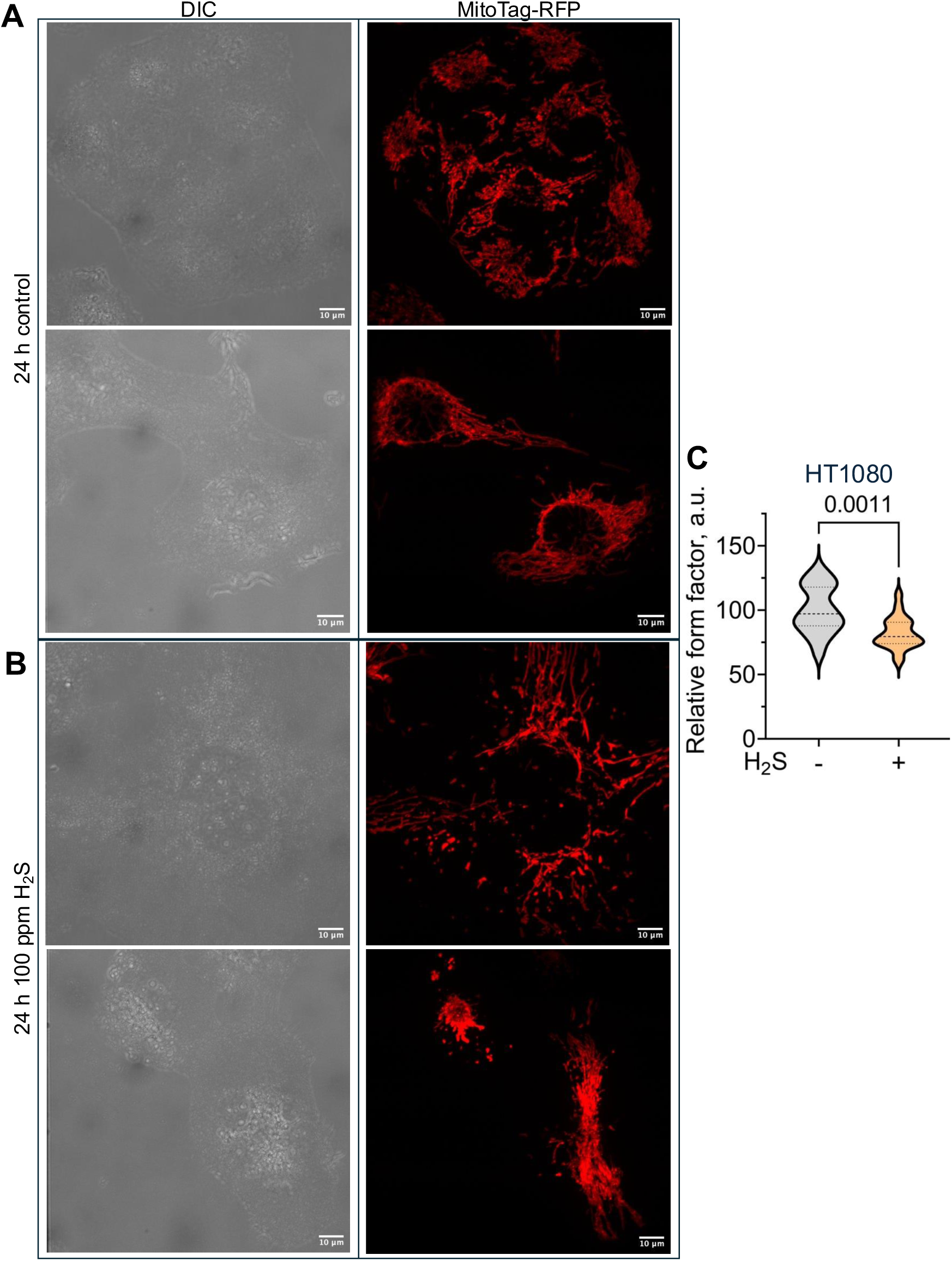
Chronic H_2_S decreases mitochondrial networking in HT1080 cells. (**A,B**) Mitochondrial morphology in control (A) versus (B) H_2_S-grown cells (100 ppm, 24 h) altered mitochondrial morphology and decreased mitochondrial networking. Two representative images are shown to illustrate variations in response to H_2_S. (**C**) Form factor analysis revealed decreased mitochondrial networking (n > 20 images).

**Figure S4.**
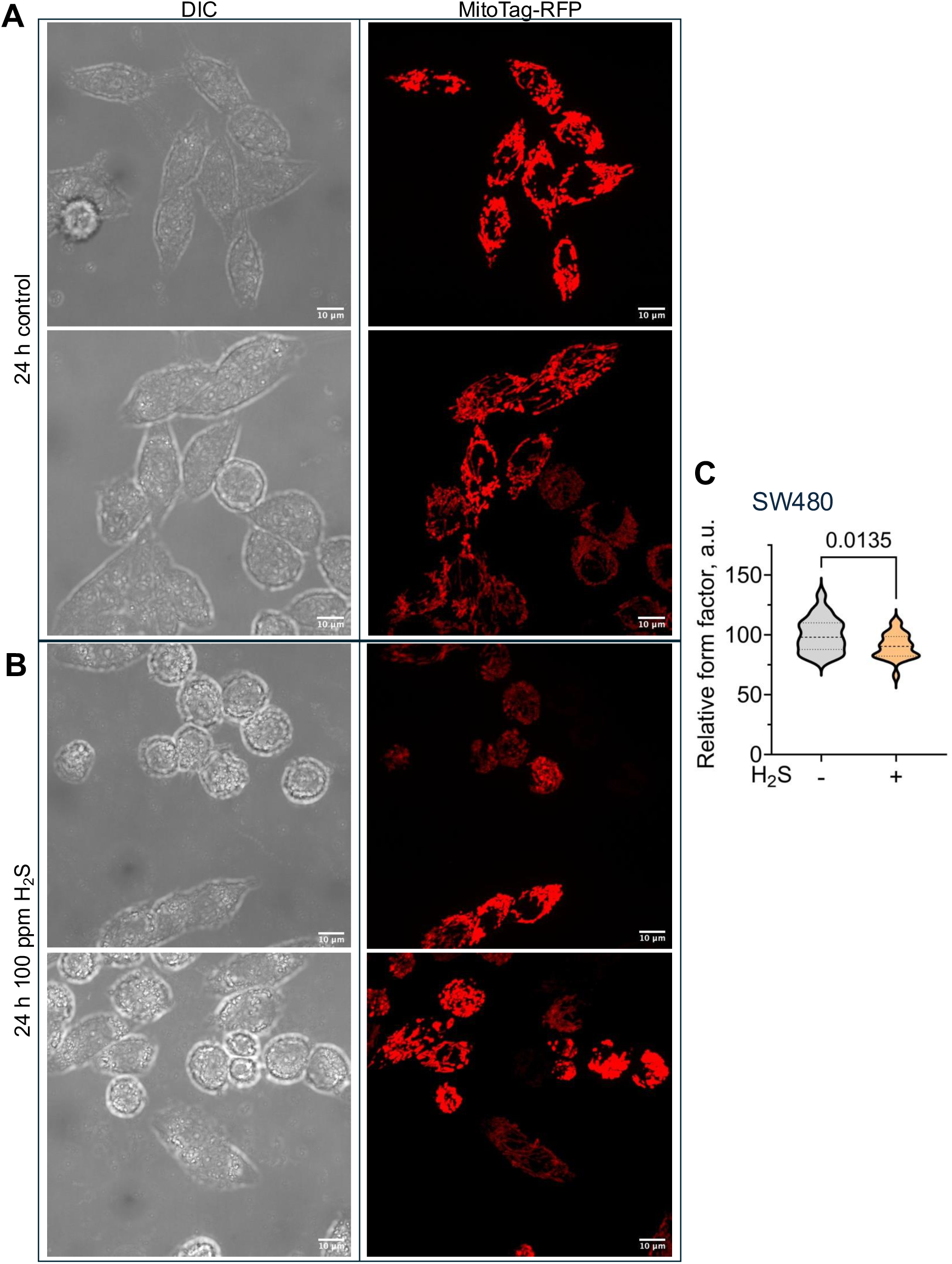
Chronic H_2_S decreases mitochondrial networking in SW480 cells. (**A,B**) Mitochondrial morphology in control (A) versus (B) H_2_S-grown cells (100 ppm, 24 h) altered mitochondrial morphology and decreased mitochondrial networking. Two representative images are shown to illustrate variations in response to H_2_S. (**C**) Form factor analysis revealed decreased mitochondrial networking (n >20 images).

**Figure S5.**
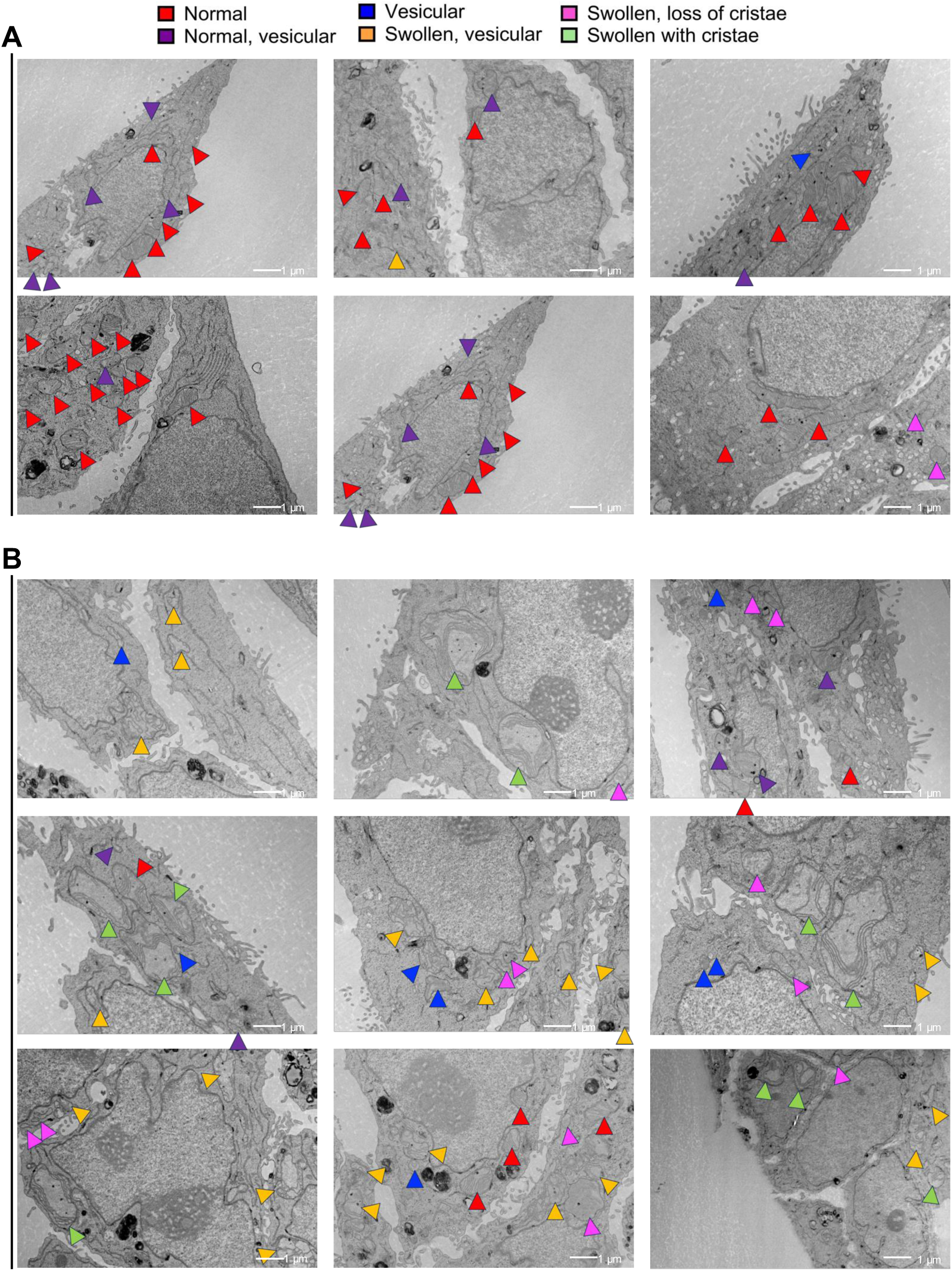
Transmission electron microscopy analysis of H_2_S effects on mitochondria. **(A,B)**. Representative images of ultrastructural changes in mitochondria in control (A) and H_2_S cultured (B, 100 ppm, 24 h) HT-29 cells (scale bar: 1 µM). Ultrastructural morphologies are annotated as described at the top.

**Figure S6.**
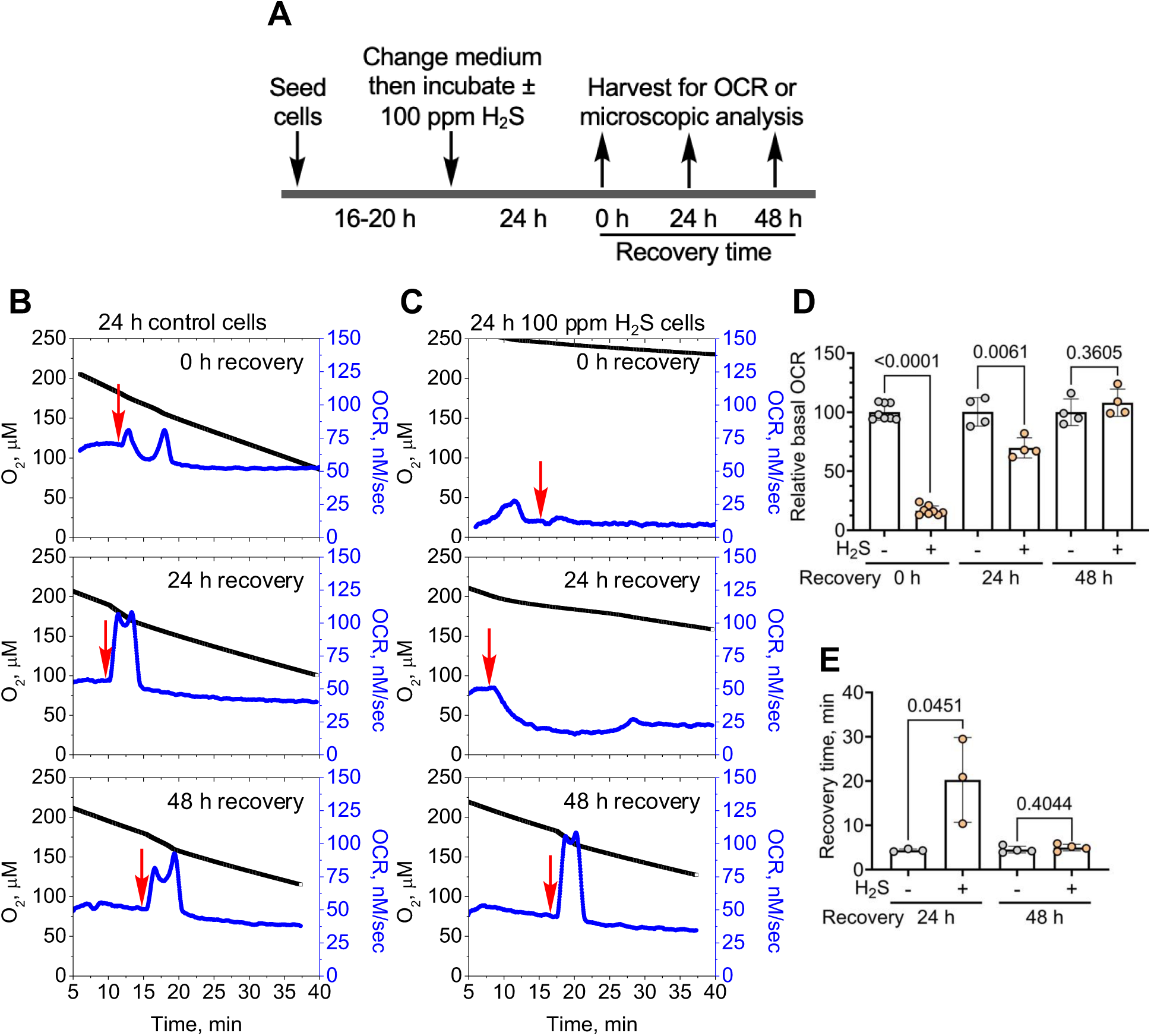
Chronic H_2_S exposure impairs respiration. **(A)** Scheme illustrating the experimental setup. The recovery times refer to the duration of culture without sulfide after an initial exposure (100 ppm H_2_S, 24 h). (**B,C**) In contrast to control cells, which respond to the addition of 20 µM H_2_S (red arrows) by increasing OCR, H_2_S-grown cells exhibit very low basal OCR and are unresponsive to exogenous sulfide (B,C *top*) but show signs of recovery 24 and 48 h after removal from the H_2_S chamber(B,C, *middle and lower*). (**D, E**) Quantitation of the basal OCR (D) and recovery time (E) data in B and C. The OCR data are representative of 3-4 independent experiments.

**Figure S7.**
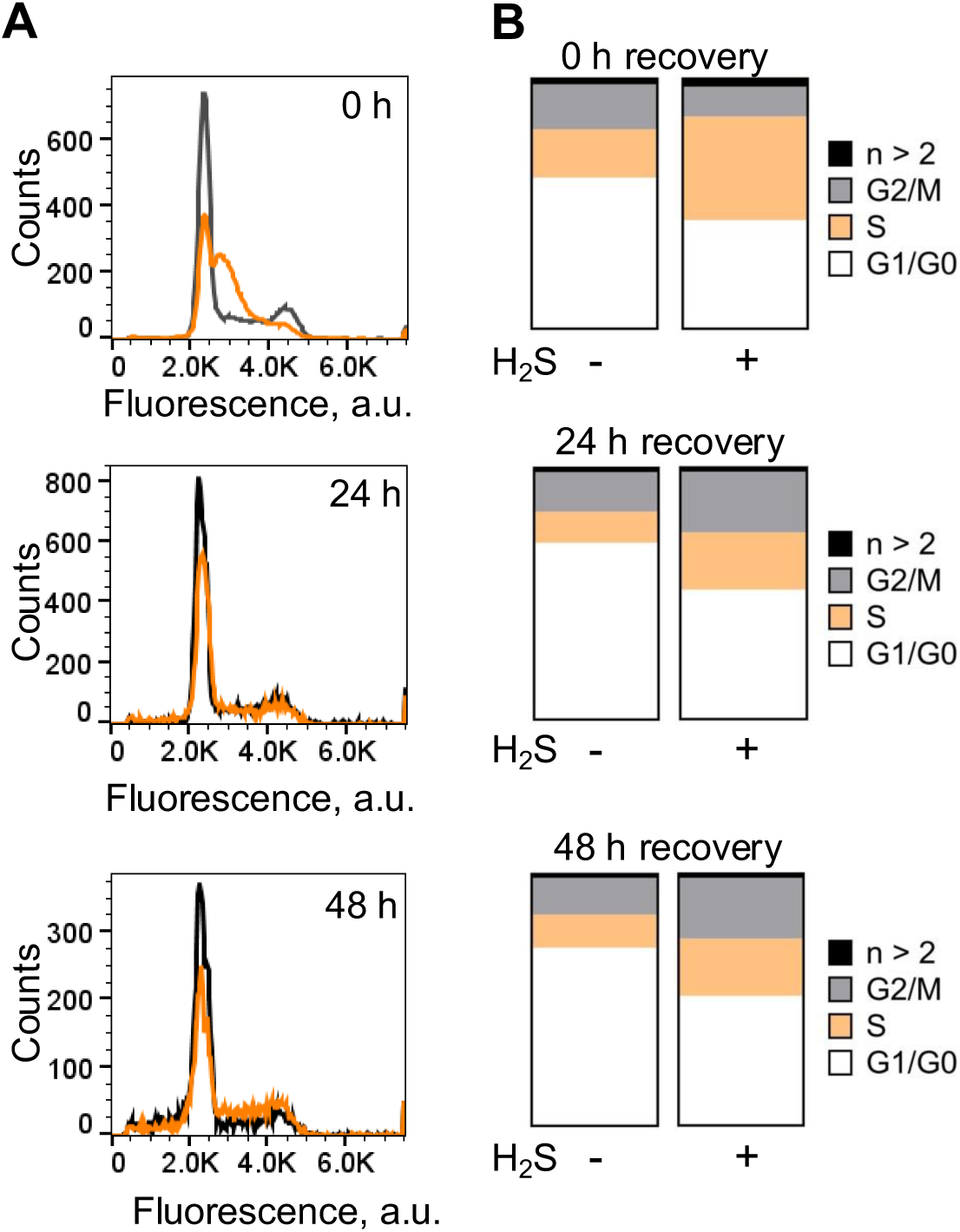
Sulfide induces S-phase arrest. (**A,B**) HT-29 were grown ± H_2_S (100 ppm, 24 h) and then allowed to recover for the next 48 h in the absence of sulfide. Flow cytometry analysis of DNA content (B) and quantitation of data (C) indicate S-phase cell-cycle arrest after 24h of sulfide exposure, which is subsequently relieved over 48 h of culture in the absence of H_2_S. The data are representative of 2 independent experiments each conducted in triplicate.

**Figure S8.**
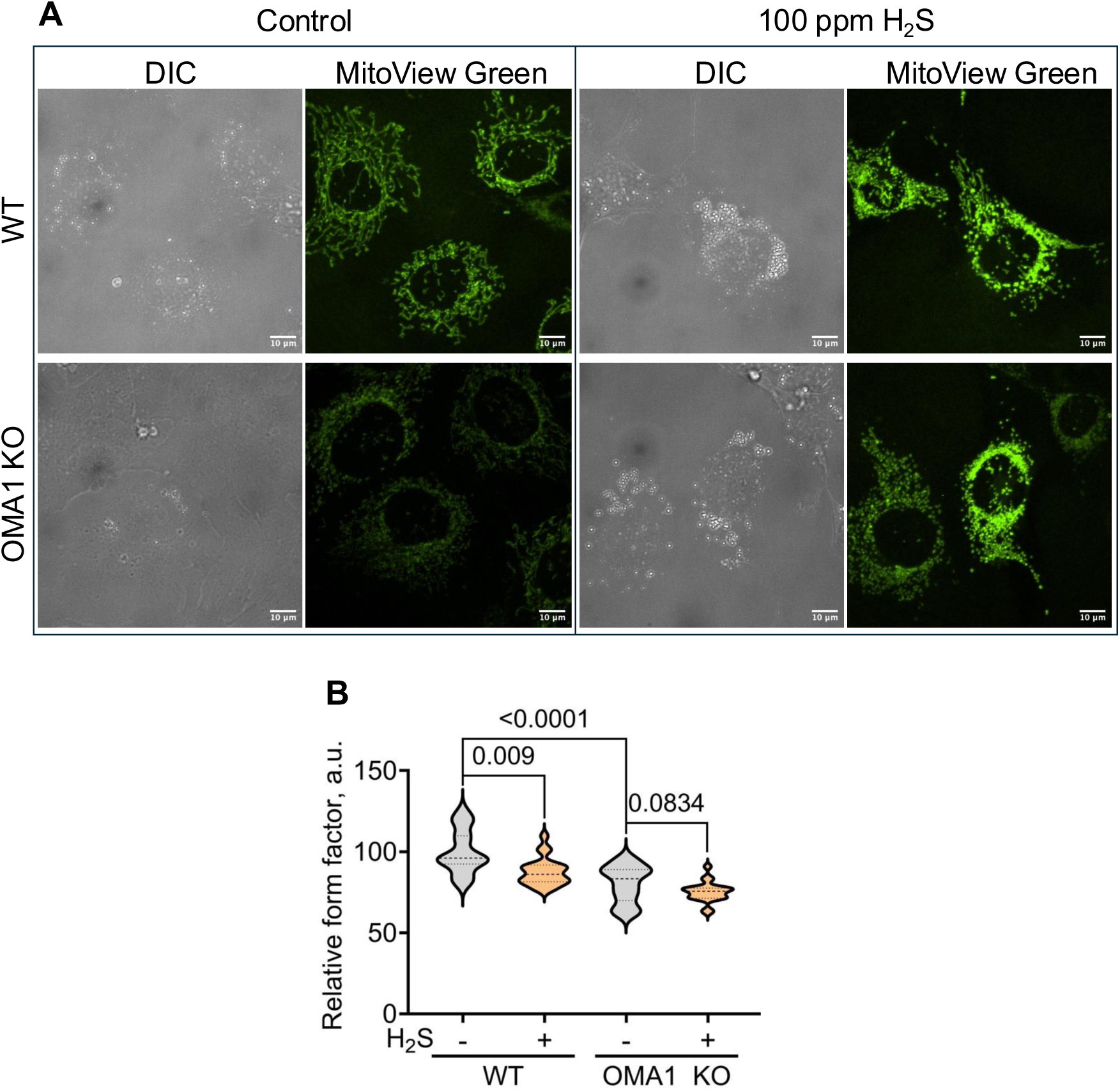
H_2_S-dependent decrease in mitochondrial networking is OMA1-dependent. (**A**) Representative microscopy images of wild-type and OMA1 KO MEF cells (cultured ± 100 ppm H_2_S, 24 h). OMA1 KO cells exhibit lower intensity for mitochondrial staining and H_2_S enhances MitoView Green staining in both samples. (**B**) Mitochondrial networks were estimated from the relative form factor of mitochondria (n = 19 to 22 images per condition from two independent experiments each conducted in duplicate). In contrast to wild-type cells, mitochondria in OMA1 KOs are less impacted by H_2_S exposure. Two-sample unpaired t test was performed for the statistical analysis in B.

**Figure S9.**
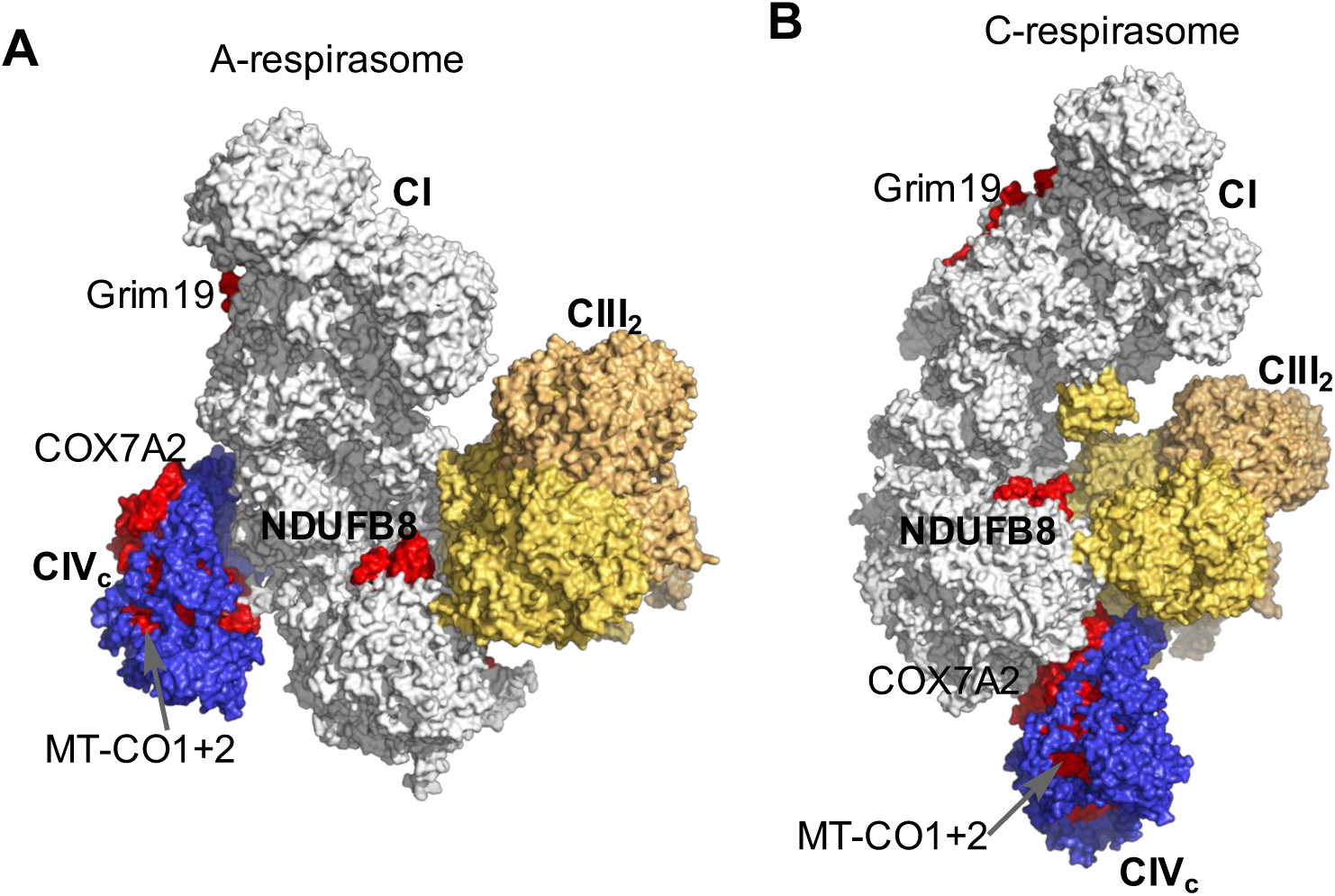
Structures of the A- and C-respirasome highlighting subunits that are affected by H_2_S. **(A, B**) Surface representation of the A- and C-type respirasomes (PDB <u>8PW5</u> and <u>8PW6</u>) depicting CI (gray), CIII dimer (CIII_2_, yellow) and CIV with COX7A2 (CIV, blue). In the A-respirasome, CIV_c_ binds exclusively to CI and COX7A2 is not predicted to contribute to the stability of the CI and CIV interaction. In the C-respirasome, CIV_c_ binds to CI and CIII_2_ and COX7A2 appears to be necessary for stabilizing CIV_c_ in this supercomplex. The CI and IV subunits that decrease in abundance in response to H_2_S are shown in red. Loss of these protein subunits, especially COX7A2 predict instability and loss of CIV from the respirasome.

